# A cancer-associated RNA polymerase III identity drives robust transcription and expression of SNAR-A noncoding RNA

**DOI:** 10.1101/2021.04.21.440535

**Authors:** Kevin Van Bortle, David P. Marciano, Qing Liu, Andrew M. Lipchik, Sanjay Gollapudi, Benjamin S. Geller, Emma Monte, Rohinton T. Kamakaka, Michael P. Snyder

## Abstract

Dysregulation of the RNA polymerase III (Pol III) transcription program, which synthesizes tRNA and other classes of small noncoding RNA critical for cell growth and proliferation, is associated with cancer and human disease. Previous studies have identified two distinct Pol III isoforms defined by the incorporation of either subunit POLR3G (RPC7α) during early development, or POLR3GL (RPC7β) in specialized tissues. Though POLR3G is re-established in cancer and immortalized cell lines, the contributions of these isoforms to transcription potential and transcription dysregulation in cancer remain poorly defined. Using an integrated Pol III genomic profiling approach in combination with *in vitro* differentiation and subunit disruption experiments, we discover that loss of subunit POLR3G is accompanied by a restricted repertoire of genes transcribed by Pol III. In particular, we observe that a specific class of small noncoding RNA, *SNAR-A*, is exquisitely sensitive to the availability of subunit POLR3G in proliferating cells. Taken further, large-scale analysis of Pol III subunit expression and downstream chromatin features identifies concomitant loss of POLR3G and *SNAR-A* activity across a multitude of differentiated primary immune cell lineages, and conversely, coordinate re-establishment of POLR3G expression and *SNAR-A* features in a variety of human cancers. These results altogether argue against strict redundancy models for subunits POLR3G and POLR3GL, and instead support a model in which Pol III identity itself functions as an important transcriptional regulatory mechanism. Upregulation of POLR3G, which is driven by MYC, identifies a subgroup of patients with unfavorable survival outcomes in specific cancers, further implicating the POLR3G-enhanced transcription repertoire as a potential disease factor.

## INTRODUCTION

The cellular division of transcriptional labor includes the RNA polymerase III (Pol III) apparatus, specially tuned for production of small, noncoding RNAs critical for translation, transcriptional and post-transcriptional regulation, and other fundamental processes^1^. Structural and biochemical studies of the Pol III complex, gene promoter architectures, and essential transcriptional regulators have extensively decoded the Pol III transcriptome and layers of transcriptional regulation^2–5^. Well-defined classes of small RNA synthesized by Pol III include tRNA and 5S ribosomal RNA (translation), 7SK and U6 small nuclear RNA (Pol II regulation and pre-mRNA splicing, respectively), RMRP and RPPH1 (processing of rRNA and tRNA, respectively), and 7SL (RNA scaffold of the signal recognition particle)^6–15^. Additional classes of Pol III-transcribed genes, whose functions remain less well understood, include vault RNA (regulation of apoptosis and autophagy)^16,17^, Y RNA (replication initiation; translation regulation)^18,19^, BC200 (translation regulation)^20^, and SNAR RNA, which interacts with RNA-binding protein ILF3^21,22^. BC200 and SNAR RNA are both aberrantly upregulated in a variety of cancer contexts, yet the underlying mechanisms and disease contributions of these RNA species remain poorly understood^23–27^.

Three distinct promoter architectures categorically classify Pol III-transcribed genes based on internal (type I, II) and external (type III) DNA sequence features and complementary transcription factor repertoires^28^. Pol III recruitment and activity is also controlled through interactions with oncogenic and tumor-suppressor signaling pathways, connecting growth-related synthesis of tRNA and other small RNAs to extracellular growth cues^29^. Specific pathways converge on the transcription factor complex, TFIIIB, as well as MAF1, a transcriptional repressor that antagonizes Pol III occupancy and activity^30–34^. However, MAF1 is a chronic repressor of Pol III activity in both proliferative and differentiated contexts, suggesting additional mechanisms may also contribute to context-specific Pol III transcription regulation at the gene level^35^. Previous genomic mapping studies of Pol III occupancy suggest some level of tissue-specific activity patterns^36–39^, and we and others have reported dynamic loss of Pol III occupancy and transcription at specific tRNA genes during cellular differentiation and exit from proliferation^40,41^. Comprehensive, high quality profiles of Pol III activity within cellular models of development are needed to better understand the level of dynamic transcription at other classes of Pol III-transcribed genes, and the mechanisms contributing to restricted Pol III activity during differentiation.

Pol III is the largest RNA polymerase protein complex and is comprised of multiple subunits that are either Pol III-specific or otherwise shared with Pol I and/or Pol II^42^. Unique components include the large subunits, POLR3A (RPC1) and POLR3B (RPC2), which constitute the catalytic DNA-binding core, POLR3D (RPC4) and POLR3E (RPC5), which establish a stably associated subcomplex involved in transcription initiation and termination, as well as POLR3C (RPC3), POLR3F (RPC6), and POLR3G (RPC7α), which together form a heterotrimeric subcomplex required for recruitment of Pol III via direct interactions with transcription factor complex, TFIIIB^43–46^. POLR3G is highly expressed during early development and subsequently turned off during differentiation^47,48^. A paralogous subunit, POLR3GL (RPC7β), is expressed at later stages of development and required for long-term survival *in vivo*^49^. Incorporation of POLR3GL into the analogous RPC3-RPC6-RPC7 heterotrimeric subcomplex altogether signifies that Pol III-transcribed genes are expressed by distinct complex identities during development.

POLR3G and POLR3GL share 46% amino acid identity and were established by a gene duplication event in the common ancestor of vertebrates^50^. Whether differences in Pol III composition defined by POLR3G or POLR3GL mechanistically drive unique transcription patterns has not been well established. However, initial mapping of POLR3G and POLR3GL, as well as functional analysis of these subunits in mice, suggests a strong level of redundancy with respect to gene occupancy and contribution to developmental potential, respectively^49,50^. Despite evidence of redundancy, knockdown of POLR3G in human cells induces changes in specific Pol III-transcribed RNA levels, and ectopic expression of POLR3G enhances the expression of several classes of Pol III-transcribed genes not otherwise affected by ectopic POLR3GL expression^48,51^. Recent Pol III structural analysis suggests that interactions between the conserved C-terminal tail regions of POLR3G and POLR3GL and the active center of Pol III may function as an autoinhibitory mechanism that precludes transcription in the absence of direct interactions with TFIIIB^52^. While potential differences in POLR3G- and POLR3GL-mediated autoinhibition require further investigation, additional structural insight suggests that subunit POLR3G (RPC7α), but not POLR3GL (RPC7β), may block the site of MAF1 interaction required for repression of Pol III activity^53^. These findings imply that POLR3G and POLR3GL may not be strictly redundant, and that complex identity may indeed exert some level of context-specific control of Pol III transcription potential.

Towards better understanding the mechanisms underlying dynamic Pol III activity and the contributions of POLR3G and POLR3GL, we established an unprecedented genomic map of nearly half of all Pol III subunits in proliferating THP-1 monocytes, and traced the effects of POLR3G loss on Pol III activity during differentiation and experimental disruption (Fig. 1a). We follow-up these experiments with large-scale analyses of Pol III identity and downstream chromatin features in primary immune cell populations and primary solid tumors. These experiments consistently provide evidence that POLR3G drives enhanced expression of SNAR-A RNA, specifically, and expansion of Pol III transcription potential, generally. Our results suggest developmental loss of POLR3G restricts the transcription potential of Pol III, supporting a nonredundant model in which subunits POLR3G and POLR3GL contribute to a dynamic transcriptional landscape, with major biological implications for human health and disease.

**Fig. 1.**
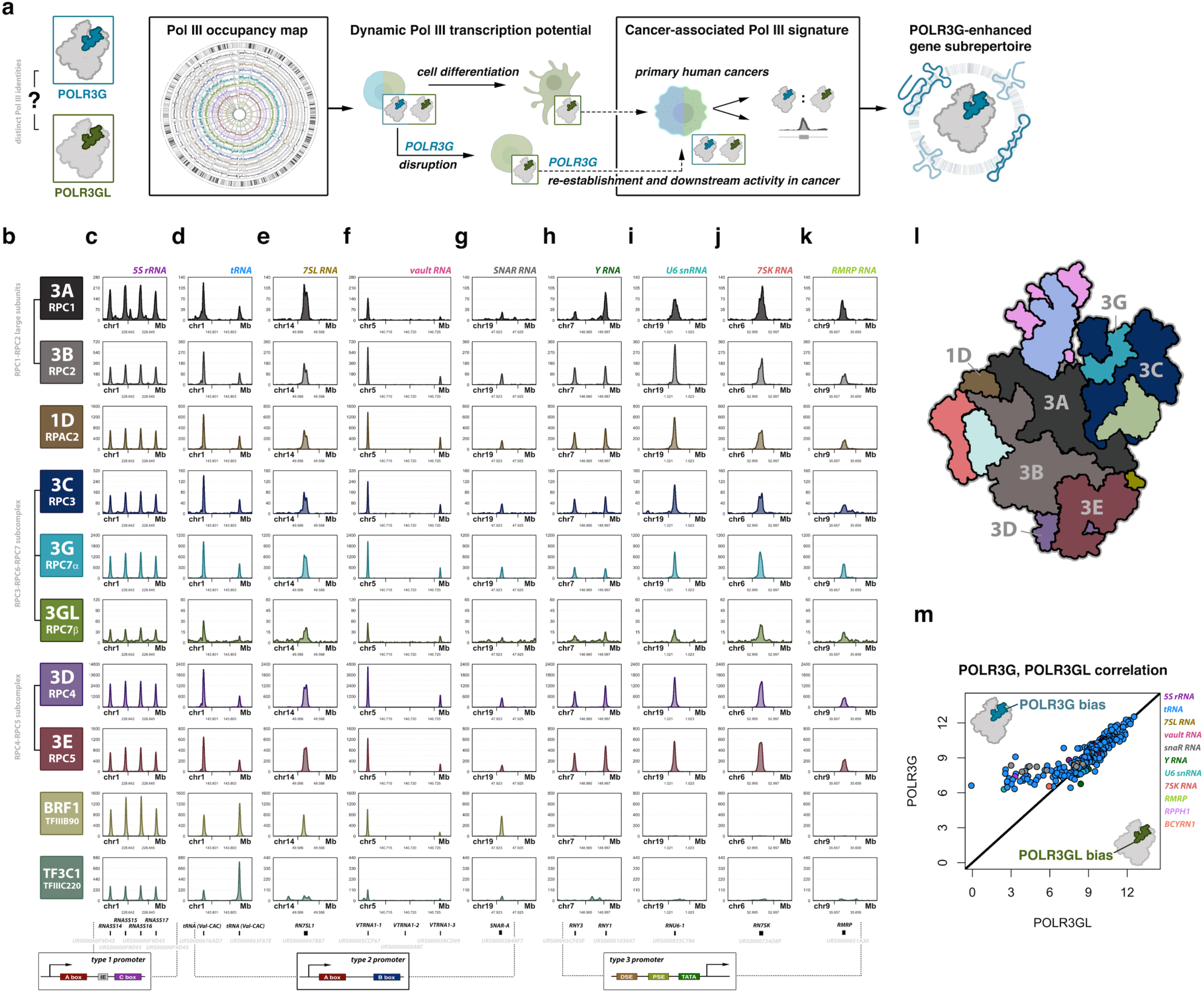
Study workflow and combinatorial atlas of human RNA Polymerase III subunits across canonical Pol III-transcribed genes. (**a**) Illustration of experimental approach tracing the effect of developmental loss of subunit POLR3G, subunit-specific disruption of POLR3G, and cancer-associated re-establishment of POLR3G and accompanying chromatin features, which collectively identify POLR3G-driven modulation of Pol III transcription potential in proliferating cells. (**b**) Pol III subunit and transcription factor legend corresponding to genome-wide maps in panels c-k, including POLR3A/RPC1 (3A); POLR3B/RPC2 (3B); POLR1D/RPAC2 (1D); POLR3C/RPC3 (3C); POLR3G/RPC7α (3G); POLR3GL/RPC7β (3GL); POLR3D/RPC4 (3D); POLR3E/RPC5 (3E); BRF1/TFIIIB90; TF3C1/TFIIIC220. Corresponding subcomplex indicated. (**c-k**) Example ChIP-seq read signals for each subunit/factor are shown in THP-1 monocytes across canonical Pol III-transcribed genes of varying promoter architecture, including ribosomal 5S rRNA genes (type 1 promoter; panel **c**), tRNA, 7SL, vault, and SNAR RNA genes (type 2 promoter, panels **d-g**, respectively), and Y, U6, 7SK, and RMRP RNA genes (type 3 promoter, panels **h-k**, respectively). Gene labels include Unique RNA Sequence (URS) identifiers assigned by (and connected to) the RNAcentral database. Bottom panel illustrates corresponding promoter architecture classification. (**l**) Cartoon schematic of the Pol III protein complex with emphasis on subunits mapped in this study (labeled) and the corresponding color code for genomic signal plots. Illustration serves as general reference guide inspired by previous structural reconstructions (Hoffmann et al., Nature 2015)^45^. (**m**) Scatterplot visualization of the correlation between normalized read densities for paralogous Pol III subunits, POLR3G and POLR3GL, at Pol III complex-occupied genes in THP-1 monocytes.

## RESULTS

### A combinatorial atlas of Pol III occupancy across canonical Pol III-transcribed genes

We used a combinatorial genomic approach to identify active Pol III-transcribed genes, relying on the extended fragment size and corresponding sequence reads of chromatin immunoprecipitation (ChIP) experiments to confidently identify Pol III complex localization. Measures of Pol III occupancy are subsequently integrated with profiles of nascent and steady-state small RNA abundance. Included in our expansive genome-wide Pol III map are the binding profiles for both large subunits, POLR3A and POLR3B, shared subunit POLR1D, multiple subunits of the RPC3-RPC6-RPC7 heterotrimer subcomplex involved in transcription initiation, including POLR3C, POLR3G, and POLR3GL, and both subunits of the RPC4-RPC5 heterodimer subcomplex involved in transcription initiation and termination, POLR3D and POLR3E (Fig. 1b; 1l). We additionally profiled the binding patterns of BRF1, a subunit of general transcription factor TFIIIB, as well as TF3C1, the largest subunit of TFIIIC (Fig. 1b). The overlap of all mapped Pol III subunits at specific genomic elements is a confident indicator of Pol III transcription and, in this respect, is a valuable resource for identifying the gene repertoire of Pol III activity within this system. Pol III and transcription factor ChIP signals were analyzed over a comprehensive list of canonical Pol III-transcribed gene coordinates using RNAcentral (https://rnacentral.org), a collection of multiple noncoding RNA annotation databases^54,55^.

In total, we identified 350 canonical Pol III-transcribed genes occupied by all mapped subunits in our cell culture system (Extended Data Fig. 1a). Inspection of individual subunit ChIP-seq signal intensities confirm occupancy across all classes of Pol III-transcribed genes, including genes encoding 5S rRNA, tRNA, 7SL, vault, SNAR, Y, U6, 7SK, and RMRP RNA (Fig. 1c-k). BRF1 and TF3C1 occupancies are restricted to type 1 and type 2 promoter architectures but absent at type 3 gene promoters, consistent with the established mechanisms and transcription factor repertoires underlying Pol III transcription initiation across genes with distinct internal and upstream promoter sequence features^28^. Overall, maps of Pol III subunit occupancy are broadly consistent across individual Pol III subunits, with specific genes featuring high levels of signal intensity across each subunit, and other genes featuring moderate to low signal intensity for all mapped subunits. Rank normalization of Pol III and transcription factor signal intensities illustrate this relationship, with high occupancy genes featuring strong signals for all subunits (Extended Data Fig. 1a). Individual Pol III subunit occupancies significantly correlate across all comparison groups, as well as with measures of chromatin accessibility, suggesting that ATAC-seq experiments can provide a general prediction of Pol III-transcribed gene activity in the absence of Pol III ChIP experiments (Extended Data Fig. 1b-j). Overall, the agreement between individual Pol III ChIP experiments suggests that, to a large degree, each subunit accurately reflects the level of Pol III complex gene localization.

The paralogous Pol III subunits POLR3G and POLR3GL are expressed at similar levels in human THP-1 monocytes (Extended Data Fig. 1k). Comparison of POLR3G and POLR3GL binding patterns in THP-1 reveals a strong level of overlap in signal intensity across genes of all promoter architectures (Fig. 1c-k), consistent with results from previous ChIP-seq experiments in HeLa cells and mouse tissues^49,50^. Individual Pol III complexes have been shown to incorporate either POLR3G or POLR3GL, suggesting overlapping signal densities represent the ability of distinct Pol III complexes to be recruited independent of its subunit composition^50^. However, rank normalized signal intensities do suggest some level of subunit occupancy bias at a subset of genes, most notably where POLR3GL signal is weak, including several tRNA genes, *SNAR-A*, *RPPH1*, and *BCYRN1*, which encodes BC200 RNA (Fig. 1m and Extended Data Fig. 1a). Though potential biases in subunit occupancy have been reported, to date, experiments dissecting the effects of subunit loss on transcription potential at these and other genes have not been fully explored.

### Loss of Pol III complex occupancy at a subrepertoire of genes following differentiation-induced depletion of subunit POLR3G

POLR3G is highly expressed during and critical for early development as functional null mutations in POLR3G result in embryonic lethality in mice^47,49^. During stem cell differentiation, a gradual loss of POLR3G expression coincides with an increase in the expression of POLR3GL, which becomes essential for long-term survival during mouse development^48,49^. Our analysis of Pol III subunit expression levels across a diverse array of cellular contexts in humans further corroborates the developmental transition in Pol III isoform expression, with high expression of POLR3G and low expression of POLR3GL in embryonic stem cells, and conversely low expression of POLR3G and high expression of POLR3GL across diverse cell-types and specialized tissues (Extended Data Fig. 2a). Like THP-1, analysis of Pol III subunit gene expression levels in immortalized cell lines show co-expression of both POLR3G and POLR3GL, consistent with previous mapping studies (Extended Data Fig. 2b). These results further confirm and serve to emphasize that while proliferative and pluripotent cellular contexts rely on a distinct Pol III complex identity from that utilized during and post-differentiation, there are additional contexts in which both POLR3G and POLR3GL are available for Pol III transcription.

Exposure of THP-1 cells to phorbol 12-myristate 13-acetate (PMA) induces differentiation and entry to a quiescent cellular state^56^. The transition of suspension-growing, immature THP-1 monocytes to adherent macrophages lends a simple differentiation system for isolating pure cell populations for genomic experiments^57–59^. Following monocyte-to-macrophage differentiation, we find that expression of POLR3G is significantly downregulated, whereas POLR3GL expression is relatively unaffected (Fig. 2a). Downregulation of POLR3G expression coincides with significant depletion of POLR3G protein levels specifically after 72-hour PMA treatment (Fig. 2b). These data demonstrate a switch in Pol III isoform abundance in THP-1 macrophages that is altogether consistent with the dynamic expression patterns observed for POLR3G and POLR3GL in other developmental contexts. ChIP-seq experiments identify robust loss of POLR3G occupancy at Pol III-transcribed genes following THP-1 differentiation, confirming that changes in subunit expression lead to corresponding changes in Pol III complex identity at the DNA level. Whereas POLR3G occupancy is significantly lost across all gene classes (Fig. 2c), POLR3GL occupancy remains comparatively stable, with significant changes at only a subset of Pol III-transcribed genes, which include examples of both up- and downregulated binding (Fig. 2d). The dynamic expression of POLR3G and resulting DNA-binding profiles of POLR3G and POLR3GL show that while proliferative THP-1 monocytes utilize Pol III complexes composed of either subunit, quiescent THP-1 macrophages predominantly rely on Pol III complexes containing POLR3GL.

**Fig. 2.**
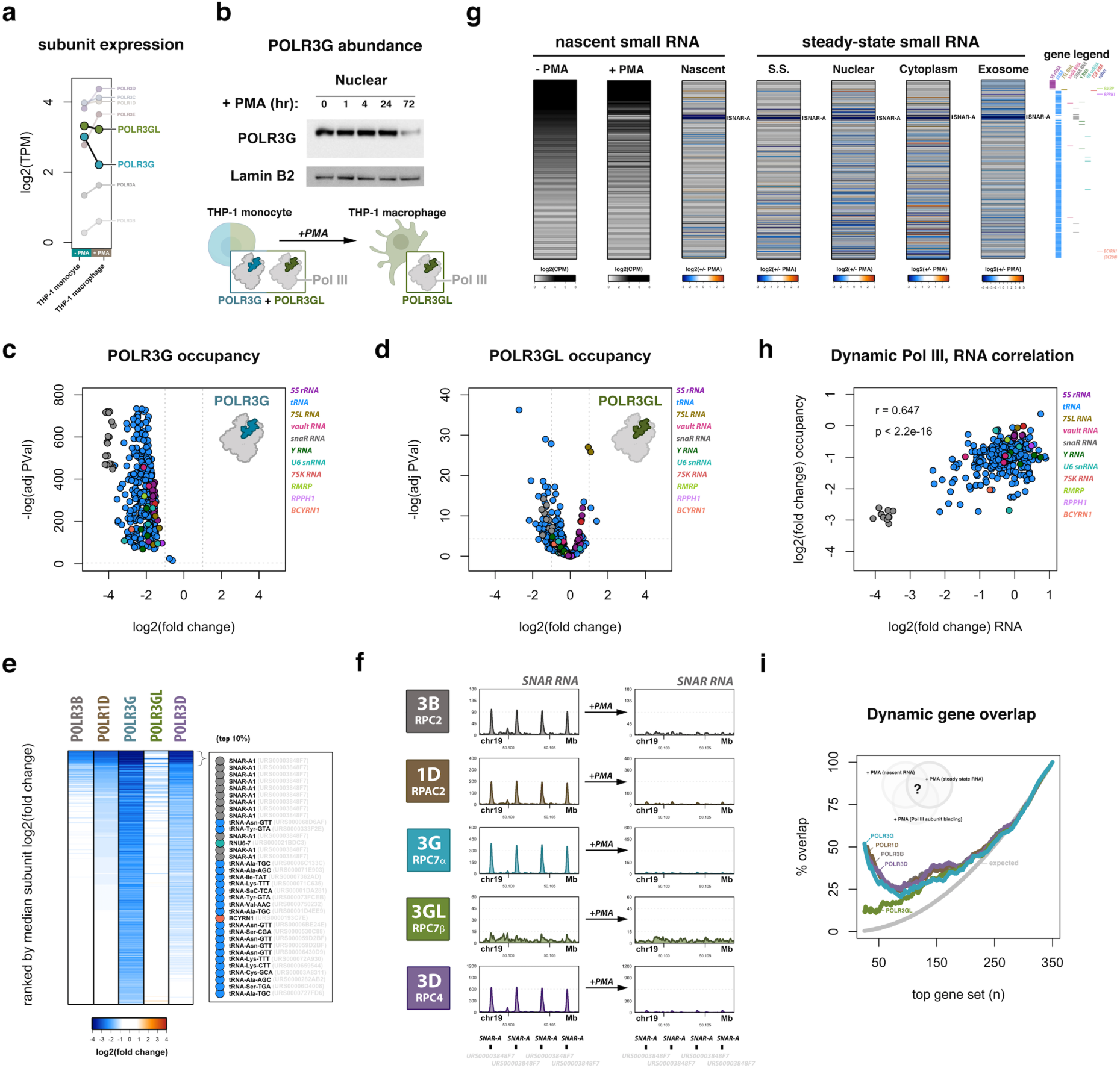
Concomitant loss of POLR3G and Pol III activity modulates cellular and exosomal small RNA repertoires during THP-1 differentiation. (**a**) Dynamic Pol III subunit expression profiles in THP-1 cells +/− 72 h PMA-induced differentiation and exit from cell proliferation. (**b**) Nuclear fractionation time course immunoblot for POLR3G and Lamin B2 protein abundance in THP-1 cells at 0, 1, 4, 24, and 72 h post PMA treatment. (**c**) Volcano plot visualization of dynamic POLR3G genomic occupancy in THP-1 cells +/− 72 h PMA-induced differentiation. Points represent individual Pol III occupied genes, gene class indicated by color legend. (**d**) Volcano plot visualization of dynamic POLR3GL genomic occupancy in THP-1 cells +/− 72 h PMA-induced differentiation. (**e**) Heatmap visualization of log2(fold change) in Pol III subunit occupancy for POLR3B, POLR1D, POLR3G, POLR3GL, and POLR3D in THP-1 cells +/− 72 h PMA. Heatmap is ordered by the median fold change across all subunits. Inset highlights the top 10% cohort of differential genes occupied by Pol III in THP-1 cells. (**f**) ChIP-seq track visualization for POLR3B, POLR1D, POLR3G, POLR3GL, and POLR3D are shown in THP-1 cells before (left) and after 72 h PMA treatment (right) across a subset of *SNAR-A* genes encoding small NF90-associated RNA. (**g**) Heatmap visualization of small RNA profiles in THP-1 monocytes, including analyses of nascent, steady-state, nuclear, cytoplasmic, and exosomal small RNA enrichment. All heatmaps are ordered by the level of nascent RNA abundance on far left heatmap. Corresponding gene legend for all heatmaps on right – rows include all canonical Pol III-transcribed genes occupied by Pol III subunits in THP-1 monocytes. From left-to-right, heatmaps visualize [1] nascent RNA abundance (- PMA) [2] nascent RNA abundance (+ PMA) [3] log2(fold change) nascent RNA (+/− PMA) [4] log2(fold change) total steady-state small RNA (+/− PMA) [5] log2(fold change) nuclear fraction steady-state small RNA (+/− PMA) [6] log2(fold change) cytoplasmic fraction steady-state small RNA (+/− PMA) [7] log2(fold change) purified exosome steady-state small RNA (+/− PMA). (**h**) Correlation between the median Pol III subunit fold change and median change in RNA abundance in THP-1 cells +/− PMA. (**i**) Moving Venn-diagram overlap analysis of the highest differential genes in both nascent and steady-state small RNA profiling experiments with differential occupancy of individual Pol III subunits.

Analogous genome-wide maps for Pol III subunits POLR3B, POLR1D, and POLR3D uncover loss of ChIP-seq signal across several classes of Pol III-transcribed genes, suggesting Pol III complex occupancy may be disrupted by the loss of POLR3G (Fig. 2e). Ranking genes by the median fold change in subunit occupancy after differentiation identifies a specific subset of Pol III-transcribed genes that exhibit substantial loss in ChIP-seq intensities for POLR3G, POLR3B, POLR1D, and POLR3D (Fig. 2e). Inspection of the top 10% of differentiation-sensitive loci reveals that genes encoding *SNAR-A*, *BCYRN1*, and multiple tRNA species are particularly depleted for Pol III localization after PMA treatment. Visual inspection of individual genes marked by diminished Pol III occupancy clearly affirms the dramatic loss of subunit localization after 72-hour PMA treatment, including at *SNAR-A* genes, tRNA genes, and *BCYRN1* which, again, exhibit comparatively weak POLR3GL ChIP-seq signal prior to treatment with PMA (Fig. 2f and Extended Data Fig. 2c-d).

### Loss of POLR3G restricts transcription and modulates cellular and exosomal small RNA repertoires in THP-1 cells

To characterize the transcriptional and post-transcriptional signatures of THP-1 differentiation, we integrated both nascent and steady-state small RNA profiles in THP-1 monocytes and macrophages. In addition, nuclear and cytoplasmic subcellular fractionation and purification of exosomes were performed to identify compartmental enrichment and corresponding changes in small RNA levels. Visualization of nascent small RNA abundance in THP-1 monocytes identifies comparatively high expression of 5S ribosomal RNA, RMRP, 7SL, 7SK, and RPPH1, moderate-to-high expression of vault, SNAR-A, U6 spliceosomal, and Y RNA, low expression of BC200, and a wide range in the expression levels for Pol III-occupied tRNA genes (Fig. 2g). Nuclear and cytoplasmic fractionation experiments appropriately identify nuclear enrichment of RMRP and U6 spliceosomal RNA, and cytoplasmic enrichment of 7SL, SNAR-A, and most tRNA species (Extended Data Fig. 2e-g). We identify Y RNA as a dominant component of exosomal small RNA profiles in THP-1 as similarly reported in other contexts (Extended Data Fig. 2g)^60,61^. Interestingly, tRNA sequencing reads mapping to tRNA-Lys and tRNA-Gly, which have been recently described to be elevated in plasma exosomes of cancer patients, are also significantly enriched in THP-1 monocyte exosomes (Extended Data Fig. 2g)^62^.

Visualization of nascent small RNA abundance in THP-1 macrophages identifies significant loss in transcription for a subset of Pol III-transcribed genes after 72-hour PMA treatment (Fig. 2g). Several changes in nascent transcription are similarly captured in steady-state and cellular fractionation small RNA-seq experiments, suggesting that changes in transcription influence the cellular availability of specific noncoding RNA after differentiation. Consistent with the dramatic loss of Pol III occupancy observed at the *SNAR-A* gene clusters on chromosome 19, the most significant change in nascent and steady-state RNA abundance after PMA treatment is attributed to SNAR-A RNA. The significant loss of SNAR-A abundance is observed across all fractionation experiments, including purified exosomes, suggesting that dynamic Pol III activity at these genes is broadly influencing both the cellular and extracellular small RNA repertoires in THP-1 cells (Fig. 2g). While fractionation small RNA-seq experiments show significant enrichment for SNAR-A RNA in the cytoplasm, SNAR-A abundance is not particularly enriched within the exosomes of THP-1 monocytes compared to total cellular RNA levels (Extended Data Fig. 2g). While this is the first study to detect dynamic, exosomal SNAR-A RNA, whether SNAR-A are actively packaged or serve any specific role in circulating extracellular vesicles remains unknown.

Overall, the dynamic RNA patterns observed during THP-1 differentiation significantly correlate with changes in Pol III occupancy, consistent with the expectation that loss of Pol III recruitment and/or occupancy should lead to changes in transcription status (Fig. 2h). Whereas the most highly differential occupancy profiles for POLR3B, POLR1D, POLR3D, and POLR3G significantly overlap genes with the most substantial changes in nascent and steady-state RNA abundance, this effect is comparatively weaker for POL3GL (Fig. 2i). Overlap enrichment patterns are consistent with the significant loss of POLR3B, POLR1D, POLR3D, and POLR3G at a specific subset of Pol III-bound genes that are marked by weaker binding signal and differential occupancy of Pol III subunit POLR3GL (Fig. 2e). These results altogether demonstrate that monocyte-to-macrophage differentiation is accompanied by a restricted repertoire of Pol III-transcribed genes concomitant with loss of subunit POLR3G. Perhaps equally important and worth emphasizing is the observation that many Pol III-transcribed genes are only modestly sensitive or seemingly unaffected in our differentiation system, including genes encoding Y RNA, RMRP, 7SL, 7SK, and RPPH1. These data imply that transitions in Pol III complex identity may not substantially affect the majority of Pol III-transcribed genes, and instead that a unique subrepertoire of Pol III-transcribed genes may be particularly sensitive to the developmental loss of POLR3G.

### Coordinate loss of POLR3G and a restricted Pol III-transcribed gene repertoire across primary immune cell lineages

To test our model that loss of POLR3G restricts the transcription potential of RNA polymerase III at specific genes, we sought to identify the relationship between POLR3G expression and the level of Pol III-transcribed gene activity beyond our THP-1 monocyte-to-macrophage differentiation system. We performed an expansive analysis of Pol III subunit expression and Pol III-transcribed gene activity across hematopoietic stem cells and multiple differentiated immune cell lineages, relying on an immense resource of simultaneous gene expression and chromatin accessibility profiling experiments generated within these contexts^63,64^. ATAC-seq, which captures the accessibility of regulatory sequence elements, is also capable of identifying the level of gene accessibility at Pol III-transcribed genes, a unique feature facilitated by the short sequence length of the Pol III gene repertoire. Direct comparison of ATAC-seq signal and Pol III subunit occupancy in THP-1 identifies a significant correlation between gene accessibility and Pol III localization, confirming the utility of ATAC-seq as a general indicator of Pol III occupancy absent direct ChIP-seq experiments against Pol III subunits (Extended Data Fig. 1i-j). Thus gene accessibility analysis offers an advantageous large-scale approach for directly querying dynamic gene signatures associated with Pol III transcription, while also cutting through the complex mixture of nascent, intermediate, and processed mature small RNA species captured in steady-state RNA experiments.

Analysis of Pol III subunit expression across a multitude of primary human immune cell types again captures the differentiation-associated loss of POLR3G expression and upregulation of POLR3GL (Fig. 3a). Whereas hematopoietic stem cells and progenitor cells express both POLR3G and POLR3GL, differentiated immune effector cells, including subsets of B cells, natural killer (NK) cells, CD4+ T cells, CD8+ T cells, and γδ T cells typically express high levels of POLR3GL and limited-to-absent levels of POLR3G. Analogous expression profiling in immune cells isolated from patients with acute myeloid leukemia (AML) identifies co-expression of both POLR3G and POLR3GL in pre-leukemic hematopoietic stem cells (pHSCs), leukemia stem cells (LSC), and leukemic blast cells (Blast) (Extended Data Fig. 3a). The THP-1 cell line, which expresses both POLR3G and POLR3GL, was derived from an acute monocytic leukemia patient and likely represents an immature precursor monocyte; thus the expression profile appears to appropriately mirror that of leukemic blast and immune progenitor cells^63^. The co-expression profile of POLR3G and POLR3GL in primary hematopoietic progenitor cells also serves to confirm the relevancy of the cellular context in which both POLR3G and POLR3GL are simultaneously available for incorporation into the RNA polymerase III complex.

**Fig. 3.**
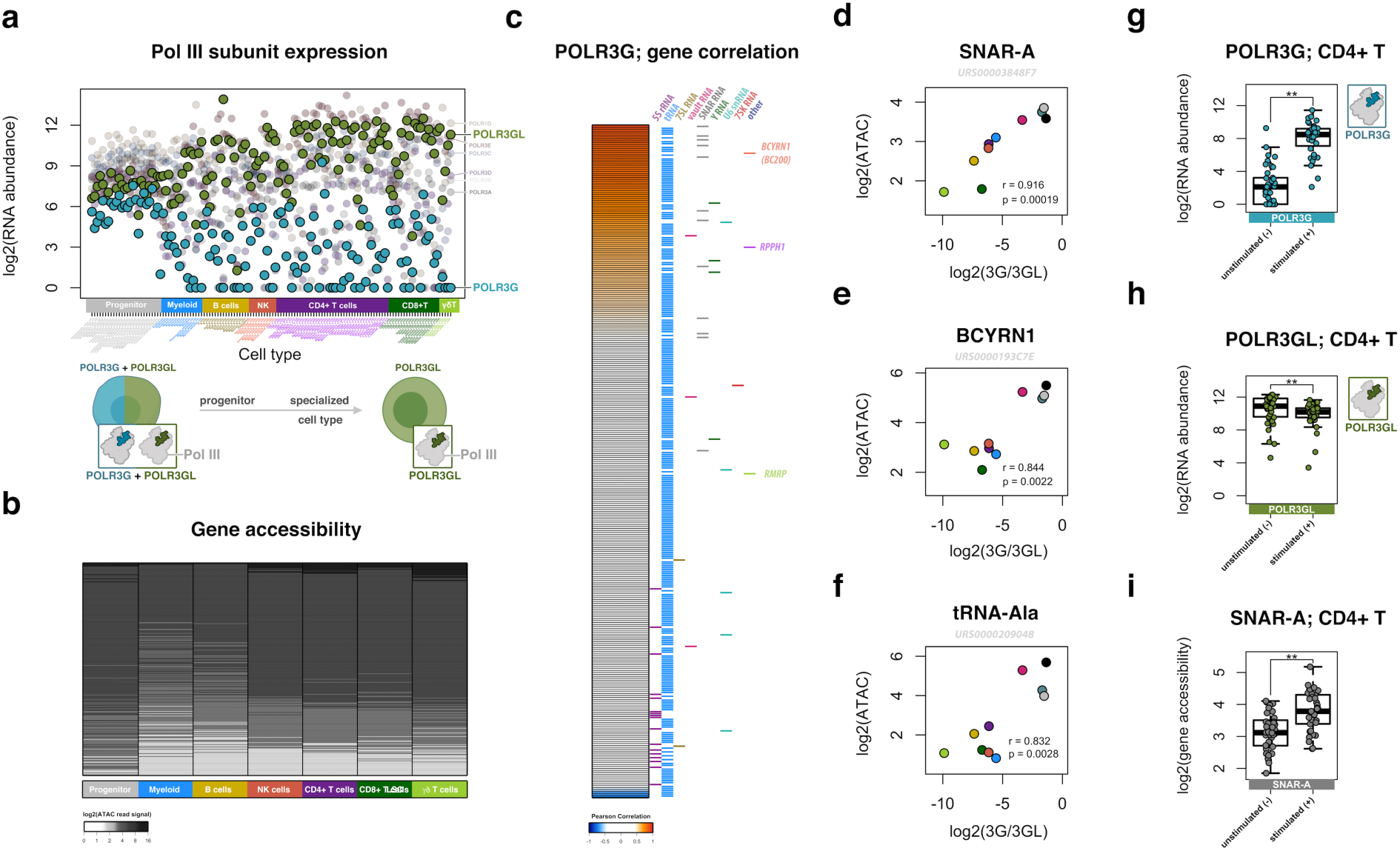
Primary immune cell differentiation is marked by loss of POLR3G expression and restricted chromatin features at a subset of genes, including *SNAR-A*. (**a**) Pol III subunit gene expression profiles across hematopoietic progenitor cells and distinct differentiation immune cell lineages, including myeloid, B cell, natural killer (NK), CD4+ T, CD8+ T, and γδ T cells, with emphasis on POLR3G and POLR3GL. (**b**) Pol III-transcribed gene accessibility as measured by ATAC-seq read signal mapping to the repertoire of Pol III occupied genes across hematopoietic progenitor cells and distinct differentiation immune cell lineages corresponding to Fig. 3a. Individual rows (genes) are ranked by median accessibility score across cell types. Loss of gene accessibility is signified by weakened ATAC-seq signal across a subset of genes in differentiated immune cell populations (**c**) Heatmap visualization of the correlation score between gene accessibility and expression ratio of POLR3G:POLR3GL. Correlation represents integration of average context-specific expression level (Fig. 3a) and average context-specific gene accessibility (Fig. 3b). Gene legend on right signifies gene type corresponding to each individual row. (**d**) Example profile of *SNAR-A* gene accessibility correlation with POLR3G:POLR3GL expression ratio in primary immune cells depicted in heatmap (Fig. 3c). (**e**) Example profile of *BCYRN1* gene accessibility correlation with POLR3G:POLR3GL expression ratio in primary immune cells. (**f**) Example profile of gene accessibility and POLR3G:POLR3GL expression ratio correlations for a specific *tRNA-Ala* (URS0000209048) gene in primary immune cells. (**g**) POLR3G expression profile in primary immune CD4+ T cell cells before and after co-stimulation with anti-CD3/CD28 Dynabeads^64^. (**h**) Analogous POLR3GL expression profile in primary immune CD4+ T cell cells before and after stimulation. (**i**) *SNAR-A* gene accessibility profile in primary immune CD4+ T cell cells before and after stimulation.

Given the significant loss of POLR3G expression within differentiated immune cell lineages, we next queried the level of gene accessibility across the Pol III transcriptome in all tested immune cell contexts. Strikingly, in comparison with the ATAC-seq profile in immune progenitor cells, all differentiated immune cell lineages exhibit a loss of gene accessibility at a specific subset of Pol III-transcribed genes (Fig. 3b). This feature is largely absent from the chromatin profile in pHSC, LSC, and Blast cells, where POLR3G expression is only modestly decreased in leukemic blast cells (Extended Data Fig. 3b). Overall, the diminishing accessibility of specific Pol III-transcribed genes concomitant with loss of POLR3G expression is broadly consistent with our model derived from experiments in THP-1, in which depletion of POLR3G leads to loss of expression at specific genes. We therefore next asked whether the subrepertoire of putative POLR3G-sensitive genes in THP-1 are similarly disrupted in our expanded profiles of immune cell differentiation.

### A subrepertoire of Pol III-transcribed genes, including *SNAR-A*, correlate with POLR3G expression bias across immune cell lineages

The measurement of both Pol III subunit expression and gene accessibility across a multitude of immune cell types allows for direct comparison of predicted gene activity with POLR3G expression. We specifically considered the ratio of POLR3G:POLR3GL expression, which accounts for the relationship between these two subunits and potential biases that arise due to higher levels of a specific subunit (Extended Data Fig. 3c). Significant correlations are observed between the chromatin signatures at a subset of Pol III-transcribed genes and the level of POLR3G expression bias in both individual primary immune samples and aggregate cell type analyses (Fig 3c-f and Extended Data Fig. 3d-h). Visualization of the correlation scores between gene accessibility and POLR3G:POLR3GL expression in aggregate cell populations again identifies a subrepertoire of Pol III-transcribed genes that appear to be modulated by POLR3G availability (Fig. 3c). Consistent with observations during THP-1 differentiation, *SNAR-A*, *BCYRN1*, and a subset of tRNA genes are among the cohort of genes with correlative chromatin features in primary immune cell populations (Fig. 3c). Direct visualization of individual genes highlights this relationship: gene accessibility is low in differentiated immune cell lineages and high in progenitor and leukemia-related contexts with high expression of POLR3G (Fig. 3d-f and Extended Data Fig. 3f-h). Importantly, the chromatin signatures at most Pol III-transcribed genes do no correlate with the dynamic identity shift from POLR3G to POLR3GL across cell populations (Fig. 3c). Visualization of *RMRP*, for example, highlights an example of Pol III-transcription that, like most Pol III-transcribed genes, appears entirely unfettered by the transition in Pol III complex identity favoring subunit POLR3GL (Extended Data Fig. 3i).

In addition to the scope of immune cell differentiation, simultaneous gene expression and chromatin accessibility profiling was also carried out in sorted primary immune cells following stimulation of individual cell types^64^. Stimulatory effects were particularly robust for B and T cells, leading to widespread changes in chromatin accessibility. We therefore considered the effect of immune cell stimulation on Pol III subunit expression and gene accessibility as well. Whereas POLR3G expression is specifically upregulated in B and T cells following robust stimulation (Fig. 3g and Extended Data Fig. 3j,k), expression of POLR3GL is comparatively unaffected by immune cell stimulation (Fig. 3h and Extended Data Fig. 3l,m). Once again, *SNAR-A* gene accessibility mirrors the level of POLR3G upregulation before and after stimulation: significant increases in *SNAR-A* chromatin features are identified in CD4+ and CD8+ T cells, where upregulation of POLR3G is most robust (Fig. 3i and Extended Data Fig 3n). *SNAR-A* gene accessibility also increases in stimulated B cells, although these changes do not reach statistical significance (Extended Data Fig. 3o). Overall, these results suggest that upregulation of POLR3G levels concomitantly re-establishes activity at *SNAR-A* genes, providing further evidence that *SNAR-A* genes represent a specialized subrepertoire of Pol III-transcribed genes enhanced by POLR3G and its corresponding Pol III complex identity for robust activity in the context of cell proliferation.

### POLR3G expression bias and downstream chromatin signatures in cancer

The re-establishment of POLR3G and *SNAR-A* chromatin features in stimulated B and T cell populations, which exit quiescence and re-enter a proliferative cellular state^64–66^, mirrors the context-specific expression of POLR3G in proliferative stem cells and upregulation in immortalized cell lines. POLR3G expression is also dysregulated in specific cancer contexts, such as transitional cell carcinoma^67^. With this in mind and, building on the putative POLR3G-sensitive repertoire in THP-1 and primary immune cells, we next sought to profile Pol III identity and downstream activity in cancer contexts. The Cancer Genome Atlas (TCGA), a database of clinical and integrated molecular signatures collected from primary human cancer tissues, provides an invaluable resource for exploring both dysregulated gene activity and chromatin features in a multitude of cancer contexts^68,69^. Analysis of POLR3G and POLR3GL expression levels in these data uncover significant increases in POLR3G expression bias in several primary solid tumors when compared to patient-matched normal tissues, including lung squamous cell carcinoma, lung adenocarcinoma, stomach adenocarcinoma, esophageal carcinoma, kidney chromophobe, cholangiocarcinoma, colorectal adenocarcinoma, bladder urothelial carcinoma, and uterine corpus endometrial carcinoma (Fig. 3a-j). In contrast, we find that POLR3G is not significantly higher in patient-matched breast, thyroid, and prostate primary cancer tissues, suggesting perhaps a more limited or less common dysregulation of POLR3G in these contexts or collected samples (Extended Data Fig. 4a-c). Nevertheless, higher levels of POLR3G expression across a multitude of cancer contexts suggests that the repertoire of POLR3G-sensitive genes identified in THP-1 and primary immune cells may also be dysregulated in these contexts. We therefore broadly explored both POLR3G levels and predictive chromatin features across Pol III-transcribed genes collected from matched primary solid tumors (Fig. 4k).

**Fig. 4.**
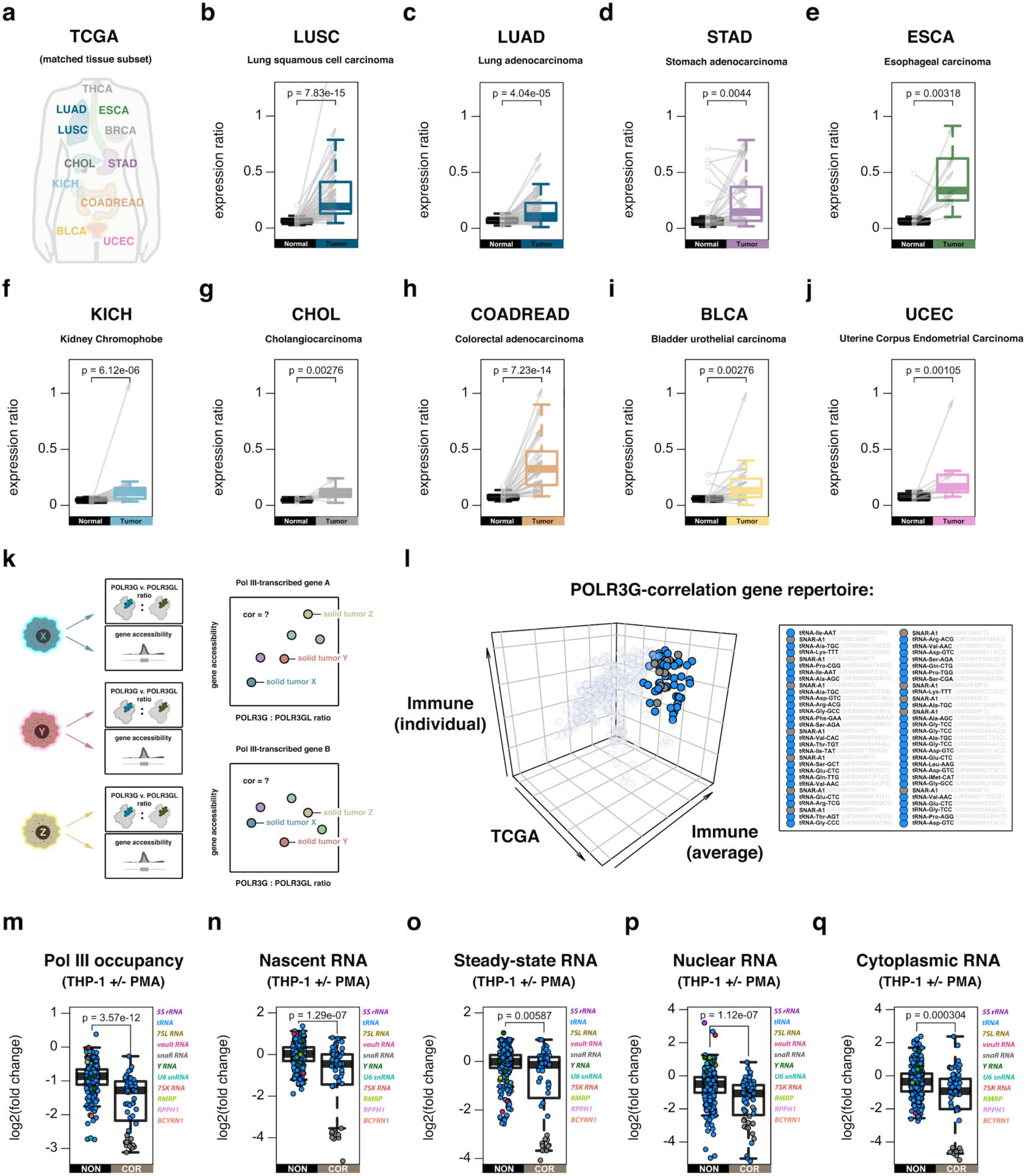
Re-establishment of POLR3G and active chromatin signatures at POLR3G-sensitive gene repertoires in human primary solid tumors. (**a**) Diagram of TCGA patient-matched normal and primary solid tumor cancer types profiled for POLR3G and POLR3GL expression levels in this study. (**b-j**) Gene expression ratios of POLR3G/POLR3GL in lung squamous cell carcinoma (**b**, n=51), lung adenocarcinoma (**c**, n=58), stomach adenocarcinoma (**d**, n=32), esophageal carcinoma (**e**, n=11), kidney chromophobe (**f**, n=25), cholangiocarcinoma (**g**, n=9), colorectal adenocarcinoma (**h**, n=32), bladder urothelial carcinoma (**i**, n=19), and uterine corpus endometrial carcinoma (**j**, n=10). Gray arrows represent individual patient-matched normal and primary solid tumors. (**k**) Illustration of workflow for integrative analysis of Pol III subunit expression and predictive chromatin features at Pol III-transcribed genes in primary solid tumors. (**l**) 3D scatterplot visualization of the subrepertoire of Pol III-transcribed genes with correlative POLR3G and predicted gene activity in The Cancer Genome Atlas (TCGA) and primary immune profiling studies. Inset notes highlighted genes with positive correlation scores in all three analyses with adjusted p value < .001 in 1 or more experiments. (**m-q**) Comparison of changes in correlative (COR) and noncorrelative (NON) subgroups related to Pol III occupancy (**m**), nascent RNA (**n**), steady-state RNA (**o**), nuclear RNA (**p**), and cytoplasmic RNA (**q**) in THP-1 cells +/− 72h PMA-induced differentiation and entry to a quiescent cellular state.

Integrated analysis of POLR3G expression biases and predictive chromatin features in individual tumor samples once again identifies significant, positive correlations between *SNAR-A* and POLR3G expression, further extending this relationship beyond THP-1 and primary immune cells into cancer contexts (Fig. 4l, Extended Data Fig. 4d). More broadly, observed cancer correlation features strongly overlap the repertoire of positive correlation features in primary immune differentiation experiments (Extended Data Fig. 4e). Inspection of Pol III-transcribed genes with positive correlation scores in both cancer and primary immune differentiation contexts identifies *SNAR-A* as well as a subset of tRNA genes (Fig. 4l). Overall, this subrepertoire of Pol III-transcribed genes, identified only through correlative chromatin features in immune cells and primary solid tumors, is more significantly downregulated during THP-1 differentiation with respect to Pol III occupancy, nascent RNA, and total, nuclear, and cytoplasmic steady-state RNA levels compared to noncorrelative genes (Fig. 4m-q). In the case of Pol III occupancy, nascent, and nuclear small RNA profiles, which more accurately reflect dynamic transcription patterns, this significance is independent of *SNAR-A* representation, further suggesting that the expression of specific tRNA genes is likely enhanced by subunit POLR3G (Extended Data Fig. 4f-j).

### Subunit-specific disruption of POLR3G induces rapid loss of SNAR-A and specific tRNA species

The Pol III-specific inhibitor, ML-60218, was discovered as an analog of small molecule inhibitors of Pol III transcription in *S. cerevisiae* with high potency in human cells^70^. The mechanism of ML-60218 has been recently characterized as POLR3G specific: ML-60218 exposure causes POLR3G depletion and a switch from POLR3G enrichment towards POLR3GL enrichment in coimmunoprecipitation experiments with other Pol III subunits^71^. Whereas depletion of POLR3GL does not affect cell proliferation, both ML-60218 exposure and depletion of POLR3G similarly trigger proliferative arrest. The overlapping effect of POLR3G depletion and ML-60218 exposure and the reported POLR3G-specific mechanism help to explain why ML-60218 fails to effectively inhibit Pol III transcription in specific contexts, such as those recently reported during preadipocyte-to-adipocyte differentiation^30^. Instead, ML-60218 is likely an effective strategy for inhibiting Pol III transcription in early developmental windows and other contexts with high POLR3G abundance. We therefore tested the effect of ML-60218 on both Pol III occupancy and small RNA levels in THP-1 monocytes to better understand the effect of POLR3G disruption on transcription within our system.

Genome-wide mapping of POLR3G and POLR3GL subunit occupancies 4 hours post-ML-60218 exposure confirms subunit-specific disruption of POLR3G localization across most Pol III-transcribed genes (Fig. 5a). Importantly, POLR3GL occupancy is not generally disrupted by ML-60218 exposure, but rather increases in ChIP-seq signal intensity at several Pol III-transcribed genes (Fig. 5b). These results are consistent with the previously characterized mechanism of ML-60218 disruption, and highlight the utility of ML-60218 as a potent tool for disrupting POLR3G activity. We therefore further leveraged ML-60218 to measure the effect of POLR3G disruption on Pol III activity in THP-1 monocytes. Small RNA profiling before and after 4-hour ML-60218 exposure identifies significant loss in RNA abundance for a specific subset of Pol III-transcribed genes, including *SNAR-A* (Fig. 5c). To better understand the temporal dynamics of ML-60218 exposure on Pol III-transcribed small RNA, we similarly profiled the RNA levels of Pol III-transcribed genes at 1, 2, and 3 hours post ML-60218 exposure. Substantial changes are observed within 2 hours of drug exposure, demonstrating a rapid effect of POLR3G disruption on the RNA abundance of a subrepertoire of Pol III-transcribed genes, including *SNAR-A* (Fig. 5d, e). Inspection of POLR3G and POLR3GL subunit occupancies at ML-60218 sensitive Pol III-transcribed genes once again illustrates the loss of POLR3G binding at *SNAR-A* and other genes with low levels of POLR3GL, with seemingly insufficient increases in POLR3GL occupancy levels to account for POLR3G disruption (Fig. 5f-i).

**Fig. 5.**
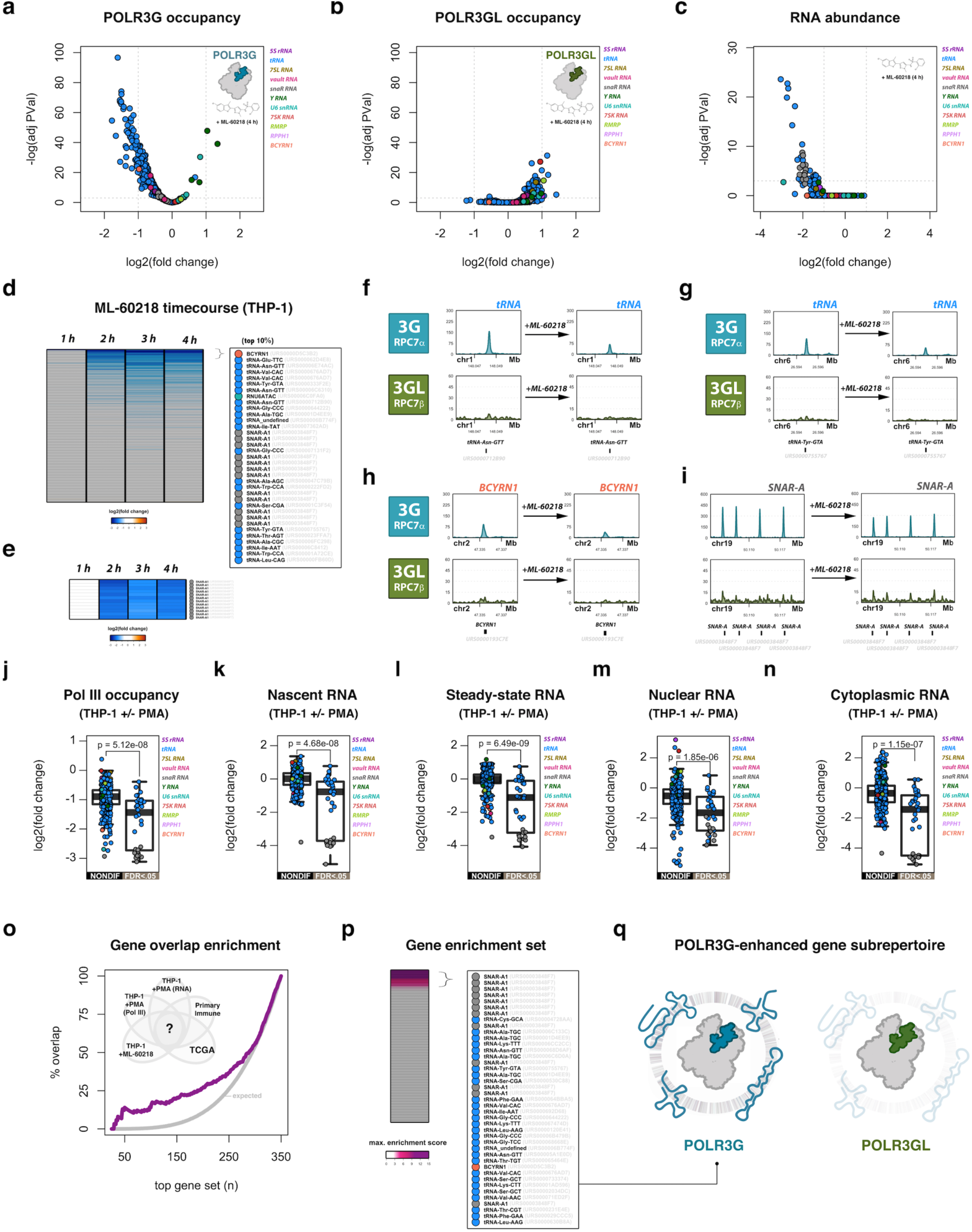
Subunit-specific drug inhibition of POLR3G induces rapid loss of POLR3G occupancy and *SNAR-A* abundance. (**a**) Volcano plot visualization of dynamic POLR3G genomic occupancy in THP-1 cells +/− 4 h exposure to Pol III inhibitor ML-60218. (**b**) Volcano plot visualization of dynamic POLR3GL genomic occupancy in THP-1 cells +/− 4 h exposure to Pol III inhibitor ML-60218. (**c**) Volcano plot visualization of small RNA abundance in THP-1 cells +/− 4 h exposure to Pol III inhibitor ML-60218. (**d**) Heatmap of log2(fold change) in small RNA abundance for Pol III-transcribed genes in THP-1 cells at 0, 1, 2, 3, and 4 h post-exposure to Pol III inhibitor ML-60218. Heatmap is ordered by the average gene log2(fold change) across the timecourse experiment. Inset highlights the top 10% differential genes. (**e**) Heatmap subset from timecourse experiment (Fig. 3d) for *SNAR-A* genes. (**f-i**) ChIP-seq track visualization for POLR3G and POLR3GL in THP-1 cells before (left) and after 4h ML-60218 exposure (right) at example tRNA genes *tRNA-Asn-GTT* (URS0000712B90) (**f**) and *tRNA-Tyr-GTA* (URS0000755767) (**g**), *BCYRN1* (**h**), and *SNAR-A* genes (**i**). (**j-n**) Comparison of changes in nondifferential (NONDIF) and significantly differential (FDR<.05) genes sensitive to ML-60218 exposure related to Pol III occupancy (**i**) and nascent RNA (**j**) in THP-1 cells +/− 72h PMA-induced differentiation. (**o**) Moving Venn-diagram overlap analysis of ML-60218 gene sensitivity 4 h post-exposure with diverse experimental results, including dynamic Pol III occupancy (THP-1 +PMA; Fig. 2), dynamic RNA abundance (THP-1 +PMA; Fig. 2), POLR3G and gene accessibility correlation in primary immune cell differentiation (aggregate cell type analysis; Fig. 3), and POLR3G and gene accessibility correlation in primary solid tumors (TCGA; Fig. 4). (**p**) Heatmap visualization of maximum enrichment scores, related to Fig. 5o, for individual Pol III-transcribed genes. Inset highlights the subrepertoire of Pol III-transcribed genes that are most likely to be enhanced by subunit POLR3G (maximum enrichment score >= 3). (**q**) Model illustration that, compared to POLR3GL, subunit POLR3G enhances the expression of specific small noncoding RNA species, including *SNAR-A*, specific tRNA genes, as well as *BCYRN1*.

Once more, Pol III-transcribed genes that are sensitive to ML-60218 exposure significantly overlap genes that are sensitive to THP-1 differentiation, independent of *SNAR-A* gene representation (Extended Data Fig. 5a-b). Significantly differential genes after 4h ML-60218 treatment are more significantly downregulated with respect to Pol III occupancy, nascent, steady-state, nuclear and cytoplasmic RNA levels during THP-1 differentiation (Fig. 5j-n), in all cases independent of *SNAR-A* gene representation (Extended Data Fig. 5c-g). Further integration of ML-60218 gene sensitivity with the dynamic Pol III and RNA signatures observed during THP-1 differentiation, and the correlative Pol III identity and chromatin features identified in primary immune and primary solid tumors, uncovers a significant overlap enrichment across these experiments (Fig. 5o). The spectrum of individual gene representation and overlap enrichment scores derived from this analysis serves to clarify the POLR3G-enhanced Pol III-transcriptome, which includes *SNAR-A* and several tRNA genes (Fig. 5p-q).

Included in the list of POLR3G-enhanced genes is *BCYRN1*, which is also among the top dynamic small RNA gene signatures identified following ML-60218-induced POLR3G disruption; however these changes do not reach statistical significance, likely due to the limited expression and read count enrichment for BC200 in steady-state small RNA profiling experiments (Fig. 5c-d). Nevertheless, loss of POLR3G occupancy at *BCYRN1* is significant, and visualization of ChIP-seq signal tracks illustrates the dynamic POLR3G binding as similarly observed during THP-1 differentiation (Fig. 5a,h). Although *BCYRN1* was not identified as a significant correlative feature with POLR3G:POLR3GL expression ratios in cancer, it was among the strongest correlative features in primary immune cells, and otherwise ranks among the top cohort with respect to concordant patterns in multiple experiments (Fig. 5o). These results suggest that, like *SNAR-A*, expression of *BCYRN1* may also be sensitive to POLR3G availability, but that additional factors may be required to increase the expression and/or post-transcriptional stability of BC200 RNA in specific contexts.

### Cancer-associated Pol III identity is driven by MYC and associated with poor survival outcomes

Previous studies have identified POLR3G-specific promoter localization of transcription factor MYC, strongly suggesting MYC regulates the expression of Pol III subunit POLR3G, but not POLR3GL^50^. In support of this finding, cistromeDB analysis of regulatory potential scores against an assemblage of transcription factor ChIP-seq experiments confirms a strong prediction score for MYC-specific regulation of the POLR3G promoter region (Fig. 6a)^72–74^. MYC is not among the predicted regulatory factors at the POLR3GL promoter, which is instead enriched for POLR2A binding, a generic signature of Pol II transcription activity and potential indicator of promoter-proximal pausing regulation of the POLR3GL gene (Fig. 6b). MYC is a well-established driver of transcriptional amplification; elevated expression and/or protein levels of MYC result in enhanced promoter occupancy and increased transcriptional output of existing gene expression programs^75^. MYC is also a central regulator of metabolic reprogramming and proliferation of stimulated B and T cell populations^76–78^, suggesting MYC likely underlies the dynamic Pol III identity signatures observed within these contexts.

**Fig. 6.**
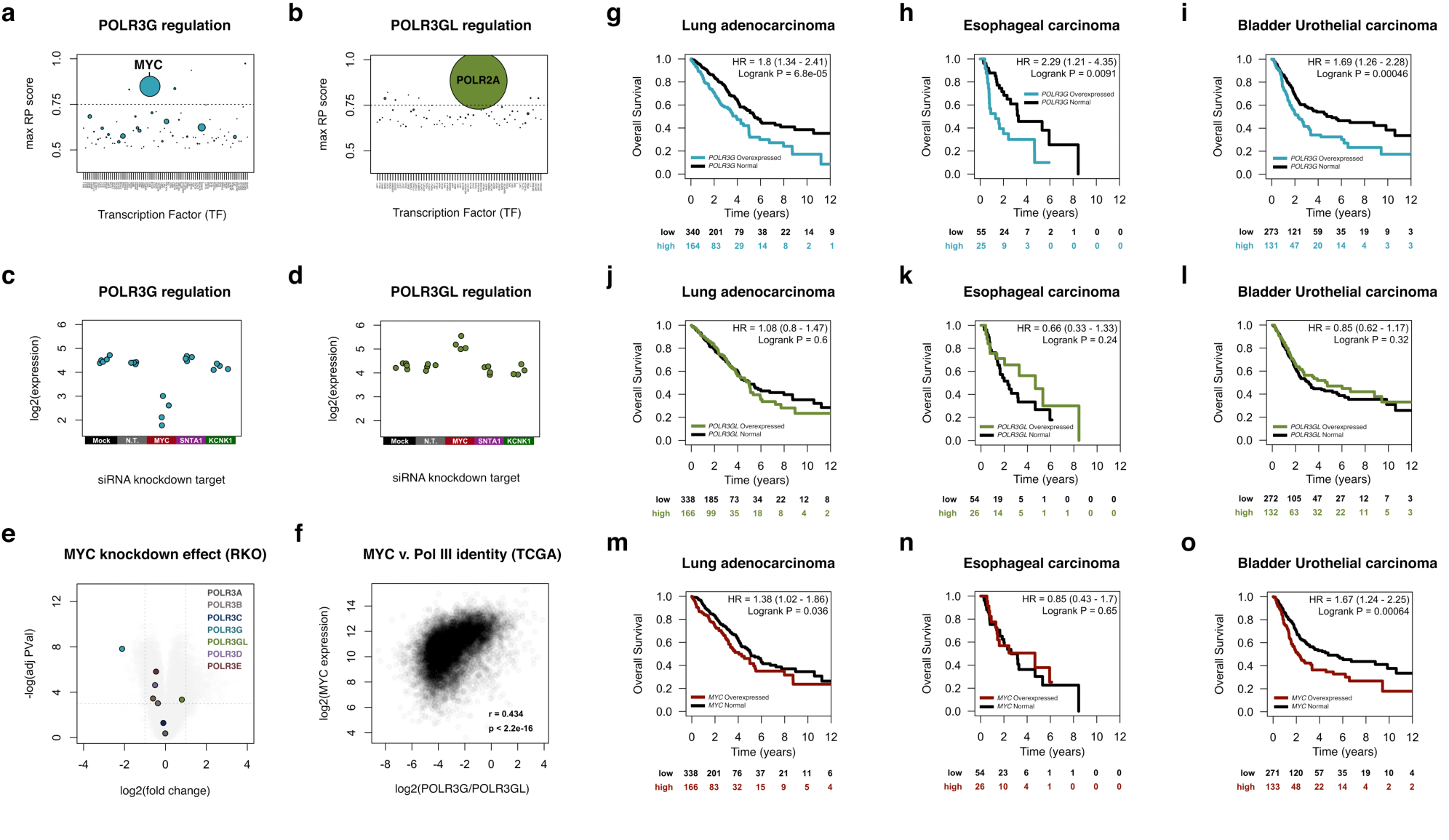
POLR3G levels are regulated by MYC and associated with poor survival outcomes in specific cancers. (**a**) Visualization of maximum regulatory potential (RP) scores for transcription factors likely to regulate POLR3G gene expression. RP scores defined by CistromeDB, which takes into account factors that bind within 1 Kb of the POLR3G transcription start site; point size corresponds to number of experiments with top RP score. (**b**) Analogous RP scores for transcription factors likely to regulate POLR3GL gene expression. (**c**) Analysis of MYC knockdown effect on POLR3G expression levels in RKO cells (Topham et al., 2015)^79^. Points represent individual biological replicates; N.T. = non-targeting. (**d**) Analysis of MYC knockdown effect on POLR3GL expression levels in RKO cells. (**e**) Volcano plot visualization of the effect of MYC knockdown on all mapped Pol III subunit expression levels in RKO cells. (**f**) Correlation analysis of MYC expression and POLR3G/POLR3GL expression ratios in TCGA primary solid tumors. (**g-h**) Kaplan-Meier analysis of overall survival of TCGA donors stratified by high POLR3G expression (top tertile) and normal POLR3G expression in lung adenocarcinoma (**g**), esophageal carcinoma (**h**), and bladder urothelial carcinoma (**i**). (**j-l**) Kaplan-Meier analysis of overall survival of TCGA donors stratified by high POLR3GL expression (top tertile) and normal POLR3GL expression in lung adenocarcinoma (**j**), esophageal carcinoma (**k**), and bladder urothelial carcinoma (**l**). (**m-o**) Kaplan-Meier analysis of overall survival of TCGA donors stratified by high MYC expression (top tertile) and normal MYC expression in lung adenocarcinoma (**m**), esophageal carcinoma (**n**), and bladder urothelial carcinoma (**o**). HR = hazard ratio risk of dying.

Robust functional knockdown experiments of MYC in proliferating RKO cells, a poorly differentiated colon carcinoma cell line, have identified a network of MYC-regulated genes with direct relevance to Pol III identity dysregulation within colon and rectal adenocarcinomas (Fig. 4h)^79^. POLR3G expression levels, which are otherwise elevated in RKO cells, are significantly diminished in response to MYC RNAi, but not in response to non-targeting siRNA or similar knockdown experiments of unrelated factors *KCNK1* and *SNTA1* (Fig. 6c). POLR3GL expression, in contrast, increases moderately following MYC knockdown, overall suggesting a significant shift in Pol III identity dependent on MYC transcription factor activity (Fig. 6d). MYC knockdown has a comparatively modest effect on the abundance of other Pol III subunits, signifying a particularly special role for MYC in modulating POLR3G abundance and downstream Pol III activity (Fig. 6e).

Building on the POLR3G and POLR3GL expression and downstream gene signatures profiled in TCGA cancer tissues, we next queried the relationship between Pol III identity and cancer-associated MYC levels and the putative effect of POLR3G dysregulation on patient survival within these contexts. MYC expression significantly correlates with POLR3G/POLR3GL expression ratios across tumor and non-tumor TCGA sample expression profiles, further supporting a model in which Pol III identity is modulated by MYC-driven transcription amplification (Fig. 6f). Kaplan-Meier analyses of overall survival in cancers often indicate concordant, adverse outcomes for elevated levels of POLR3G and MYC, but not POLR3GL (Fig. 6g-o). For example, high expression of POLR3G identifies a subgroup of patients with unfavorable survival outcomes in lung adenocarcinoma, esophageal carcinoma, bladder urothelial carcinoma, kidney renal clear cell carcinoma, uterine corpus endometrial carcinoma, and liver hepatocellular carcinoma (Fig. 6g-i and Extended Data Fig. 6a-c). In all contexts, elevated POLR3GL expression is not an indicator of adverse outcomes (Fig. 6j-l and Extended Data Fig. 6f-h). Surprisingly, while high expression of MYC is associated with adverse outcomes in specific cancers (Fig. 6m-o and Extended Data Fig. 6k-m), the hazard ratio and significance of effect is often less pronounced than POLR3G and, in certain cases, not an indicator of poor survival outcomes (e.g. esophageal cancer, Fig. 6n). These results serve to emphasize the overall significance of POLR3G and Pol III identity as a disease factor, and suggest that POLR3G-enhanced transcription may promote cancer proliferation with poor outcomes. Nevertheless, we also find that high expression of POLR3G in other specific cancer contexts, such as breast invasive carcinoma and thyroid carcinoma, where POLR3G upregulation does not appear to be a common feature, is not associated with adverse outcomes (Extended Data Fig. 6d-e), suggesting Pol III transcription dysregulation and associated outcomes are a hallmark of a multitude but subset of cancer contexts.

## DISCUSSION

Using a combination of *in vitro* differentiation and subunit disruption experiments, as well as large-scale analysis of simultaneous gene expression and chromatin signatures, our results strongly suggests that POLR3G and, consequently, Pol III identity itself, modulates transcription potential with significant implications for development and disease. In particular, we find that *SNAR-A* gene activity consistently tracks with POLR3G levels in primary immune cell populations and cancer tissues, and represents the dominant dynamic feature of both THP-1 differentiation and POLR3G disruption experiments. While this study is not the first to simultaneously map POLR3G and POLR3GL, it does not appear that the *SNAR-A* gene has been previously considered in other Pol III occupancy studies. Further, the *SNAR-A* gene is evolutionarily restricted to Hominidae^21^, and would therefore not be identified in studies of other non-Hominidae vertebrate species. The concomitant upregulation of POLR3G and re-establishment of gene accessibility at *SNAR-A* gene clusters provides compelling evidence that POLR3G may play a unique role in establishing competent and perhaps efficient transcription at these particular loci.

The observation of POLR3G-enhanced transcription of *SNAR-A* and the expanded Pol III repertoire in proliferating cells naturally demands a better understanding of the mechanism underlying POLR3G-enhanced activity. The enrichment of POLR3G signal compared to POLR3GL at *SNAR-A* and other genes suggests that Pol III complexes with POLR3G may be recruited and/or establish competent transcription at the loci with greater efficiency compared to Pol III complexes with POLR3GL incorporation. Perhaps noteworthy is the modest increase in POLR3GL signals at *SNAR-A* genes observed immediately following rapid ML-60218 disruption of POLR3G (Fig. 5i), suggesting POLR3GL incorporated Pol III complexes are not incapable of recruitment at these loci. However, the significant loss of SNAR-A RNA following POLR3G disruption again suggests that POLR3GL is likely ineffective and/or inefficient at driving transcription at these and other genes. In such a case, POLR3G would likely outcompete POLR3GL occupancy and activity, explaining the enrichment of POLR3G occupancy in THP-1 monocytes and increase in POLR3GL signal at these genes only following POLR3G disruption.

The complete loss of Pol III complex localization and lack of POLR3GL occupancy at *SNAR-A* genes following THP-1 differentiation leads us to an additional speculative model: POLR3G may localize to the *SNAR-A* gene cluster independent of the full Pol III machinery, potentially via RPC3-RPC6-RPC7 subcomplexes through interactions with TFIIIB. Recruitment might conceivably establish an accessible chromatin landscape that facilitates subsequent recruitment of Pol III for transcription. Potential supporting evidence for this model includes the comparatively moderate disruption of POLR3G occupancy at *SNAR-A* genes after 4 h ML-60218 exposure (Fig. 5a,i), despite strong disruption of SNAR-A RNA levels at the same time point (Fig. 5c-e). One interpretation of this result is that ML-60218 disrupts Pol III:POLR3G interaction but does not disrupt POLR3G-independent gene localization. A second interpretation is that, following ML-60218 disruption, POLR3GL more effectively outcompetes POLR3G at genes in which efficient transcription can be established. Other potential mechanisms include the possibility, based on recent structural data, that POLR3G rescues the activity of specific genes from MAF1-mediated repression, perhaps suggesting unique biases or efficiencies of MAF1 repression. However, we do not observe any apparent enrichment of MAF1 localization at or proximal to *SNAR-A* gene clusters that would explain potential gene bias before or following THP-1 differentiation.

The upregulation of both POLR3G and SNAR-A RNA in human cancers and adverse outcomes associated with high POLR3G expression adds further context that suggests POLR3G-driven transcription represents a disease factor. While studies of SNAR-A RNA are currently limited, SNAR-A has been shown to be upregulated in hepatocellular and ovarian carcinoma and HER2-positive breast cancer cells and, like POLR3G, immortalized cell lines^25–27^. Both POLR3G and SNAR-A promote cell proliferation, though the mechanisms for each remain poorly defined. It is expected that SNAR-A most likely functions through ILF3, the RNA-binding protein to which it interacts and to which it attributes its name^22,80^. ILF3 also promotes cell proliferation and, like POLR3G, is an unfavorable prognostic indicator across multiple cancers^81^. Future experiments dissecting the function of SNAR-A and its potential modulation of ILF3 activity will help to understand the consequence of this newly identified POLR3G-enhanced transcription program.

## METHODS

### Data acquisition

Previously reported THP-1 RNA-seq, ATAC-seq, and nascent small RNA sequencing files were obtained from Gene Expression Omnibus (GEO) series GSE96800. ATAC-seq and gene expression data derived from primary immune progenitor and primary differentiated immune cell populations were obtained from GEO series GSE74912 and GSE118119, respectively. NIH Roadmap Epigenomics gene expression profiles related to immortalized cell line (ENCODE) and multi-tissue Pol III subunit levels (Extended Data Fig. 2a,b) were extracted from the “57epigenomes.RPKM.pc” file obtained from: https://egg2.wustl.edu/roadmap/web_portal//processed_data.html#RNAseq_uni_proc. The Cancer Genome Atlas (TCGA) primary solid tumor Pol III subunit expression levels were extracted across all disease cohorts from the Broad Institute TCGA Genome Data Analysis Center (GDAC) Firehose mRNASeq Level 3 RSEM gene normalized data files at https://gdac.broadinstitute.org. Primary solid tumor ATAC-seq alignment bam files were retrieved from the Genomic Data Commons Data Portal (https://portal.gdc.cancer.gov). Clinical patient survival outcomes and Kaplan-Meier survival probability data were retrieved from the Kaplan-Meier Plotter Pan-cancer RNA-seq analysis platform with expression of POLR3G, POLR3GL, or MYC patient split set to upper tertile: (https://kmplot.com/analysis/index.php?p=service&cancer=pancancer_rnaseq). Model cell-type illustrations were created using BioRender.com.

### THP-1 cell culture

THP-1 cells were obtained from ATCC (Lot # 62454382) and grown for multiple passages in T-75 flasks between 2-8 x 10^5^ cells/mL in growth medium containing RPMI-1640 (Corning), 10% fetal bovine serum, and 1% penicillin streptomycin. For differentiation of THP-1 cells, non-adherent cells were diluted to 2 x 10^5^ cells/mL and grown overnight, and a final concentration of 100 nM PMA was added the following morning. THP-1 derived macrophages were collected after 72-hour exposure to PMA by aspirating media and any non-adherent cells, and incubating adherent cells with TrypLE (ThermoFisher) for 15 minutes followed by cell wash in phosphate buffered saline (PBS) buffer. RNA polymerase III inhibitor ML-60218 (557403-10MG) (SigmaAldrich) was added to a final concentration of 27 μM and cells collected at 1, 2, 3, and 4 hours post-exposure followed by cell wash in PBS buffer.

### Chromatin immunoprecipitation (ChIP)

Equal numbers of THP-1 monocytes, THP-1 derived macrophages, and ML-60218 treated monocytes were collected (∼10 million cells per ChIP experiment) and resuspended in growth media at 1 x 10^6^ cells/mL and cross-linked with rotation at room temperature in 1% formaldehyde for 10 minutes. Cross-linking was quenched with the addition of 200 mM glycine and an additional 5 minutes of rotation at room temperature. Cross-linked cells were then spun down and resuspended in 1x RIPA lysis buffer, followed by chromatin shearing via sonication (3 cycles using a Branson sonicator: 30 seconds on, 60 seconds off; 20 additional cycles on a Bioruptor sonicator: 30 seconds on, 30 seconds off). Individual ChIP experiments were performed on pre-cleared chromatin using antibody-coupled ChIP grade Protein G magnetic beads (Cell Signaling Technology). POLR3A antibody was obtained from Abcam (ab96328 lot#GR318563). POLR3B antibody was obtained from Bethyl (A301-855A). POLR1D antibody was obtained from Bethyl (A304-847A). POLR3C antibody was obtained from Bethyl (A303-063A). POLR3G antibody was obtained from Invitrogen (PA5-51120 lot#UG2803044). POLR3GL antibody was obtained from Novus Biologicals (NBP1-79826). POLR3D antibody was obtained from Bethyl (A302-295A). POLR3E antibody was obtained from Bethyl (A303-708A). BRF1 antibody was obtained from Abcam (ab74221). GTF3C1 antibody was obtained from SigmaAldrich (PLA0180). IgG antibody was obtained from ThermoFisher Scientific (NeoMarkers NC-100-P0). 5 ug of antibody per ChIP was coupled to 18 uL of beads and rotated overnight with sheared chromatin at 4° C. Beads were then washed 5x in ChIP wash buffer (Santa Cruz), 1x in TE, and chromatin eluted in TE + 1% SDS. Cross-linking was then reversed by incubation at 65° C overnight, followed by digestion of RNA (30 min RNase incubation at 37° C) and digestion of protein (30 min proteinase K incubation at 45° C). ChIP DNA was then purified on a minElute column (Qiagen), followed by DNA library preparation (NEBNext Ultra II DNA Library Prep Kit for Illumina) and size selection of 350-550 bp fragments via gel extraction (Qiagen).

### ChIP-seq analysis

For each individual factor-specific ChIP-seq experiment, biological replicates and experimental conditions (-PMA/ML-60218; +PMA; +ML-60218) were sequenced together on an Illumina HiSeq4000 (paired-end, 100 bp). Sequencing was performed by the Genome Sequencing Service Center by Stanford Center for Genomics and Personalized Medicine. Sequencing reads were trimmed using trim galore v.0.4.0 prior to downstream sequence alignment and analyses. Trimmed paired-end ChIP sequencing reads were mapped to the GRCh38 genome using bowtie version 2.2.4 with settings “bowtie2 -t –sensitive -x” ^82^. Read counts were extracted over a comprehensive RNAcentral database annotation of noncoding RNAs ^54,55^. Gene annotation set and corresponding GRCh38 coordinates were downloaded from the contemporaneous RNAcentral database (release version13):ftp://ftp.ebi.ac.uk/pub/databases/RNAcentral/releases/13.0/genome_coordinates/bed/h omo_sapiens.GRCh38.bed.gz. RNAcentral coordinate IDs were mapped to Ensembl, Gencode, GtRNAdb, HGNC, Lncbase, Lncbook, Lncipedia, Lndcrnadb, miRbase, noncode, Refseq, and Rfam. To remove duplicate coordinate gene entries derived from multiple databases, genes with start and end coordinates within 50 bp of each other were merged into singular coordinate entries. Gene-specific raw ChIP-seq signal counts were extracted and include the flanking 150 bp for each RNAcentral gene coordinate entry. Differential count statistics corresponding to Pol III subunit occupancy changes were derived from the exactTest function of the edgeR package for differential expression analysis ^83,84^ over the entire RNAcentral annotation set. All downstream analyses subset on canonical Pol III-transcribed genes, including tRNA, 5S rRNA, U6, 7SK, 7SL, SNAR, Y, vault, RPPH1, RMRP, BCYRN1. In total, 11,526 potential Pol III-transcribed genes, including numerous pseudogenes, were allowed for initial analysis of Pol III occupancy. After ranked assessment of Pol III subunit co-localization (Extended Data Fig. 1a), 350 unique Pol III-transcribed genes were considered to be confidently bound by the Pol III complex and likely to be transcriptionally active and considered for downstream integrated analysis with small RNA abundance. ChIP-seq signal track data were generated from post-filtering read alignment bam files using the deeptools bamCompare tool ^85,86^. Signal track visualization plots were generated using the Sushi package for genomic visualization^87^.

### Total, nuclear, cytoplasmic, and exosomal small RNA purification

Total steady-state small RNA was purified from equal numbers (∼2 million cells per small RNA-seq experiment) of THP-1 monocytes +/− ML-60218 exposure and THP-1 macrophages (72 h PMA) using the mirVana miRNA isolation kit according to instructions (AM1560, Invitrogen). Nuclear and cytoplasmic THP-1 fractions were isolated using the NE-PER Nuclear and Cytoplasmic Extraction Reagents (ThermoFisher) according to instructions, with 2 additional nuclear washes prior to nuclear extraction. Fractionation purities were assessed by β-tubulin and Lamin B2 contamination (Extended Data Fig. 2e). THP-1 monocyte and THP-1 macrophage exosomes were isolated by differential centrifugation (10 min x 300 g, 10 min x 2000 g, 30 min x 10,000 g, 70min x 100,000 g all at 4°C), washed and resuspended in PBS for analysis. Exosome particle size distributions were assessed by nanoparticle tracking analysis (NanoSight) (Extended Data Fig. 2f). Small RNA was immediately isolated from subcellular fractionation and exosomal purification experiments using the mirVana miRNA isolation kit according to instructions. Small RNA libraries were generated using the NEBnext small RNA library preparation kit according to instructions (E7580S, New England BioLabs).

### Small RNA-seq analysis

For analysis of Pol III-transcribed small RNA abundance, we mapped nascent and steady-state small RNA reads to the entire genome space (GRCh38) to avoid false positive signal arising from sequence reads unrelated to Pol III-transcribed genes that might otherwise occur using a limited reference set^88^. Sequencing reads were trimmed using trim galore v.0.4.0 prior to downstream sequence alignment and analyses. By default, multi-mapping reads were reported as a singular best alignment. Read counts were extracted over a comprehensive RNAcentral database annotation of noncoding RNAs^54,55^. Gene annotation set and corresponding GRCh38 coordinates were downloaded from the contemporaneous RNAcentral database (release version13): ftp://ftp.ebi.ac.uk/pub/databases/RNAcentral/releases/13.0/genome_coordinates/bed/homo_sapie ns.GRCh38.bed.gz. RNAcentral coordinate IDs were mapped to Ensembl, Gencode, GtRNAdb, HGNC, Lncbase, Lncbook, Lncipedia, Lndcrnadb, miRbase, noncode, Refseq, and Rfam. To remove duplicate coordinate gene entries derived from multiple databases, genes with start and end coordinates within 50 bp of each other were merged into singular coordinate entries. Gene-specific raw small RNA signal counts were extracted and include the flanking 25 bp for each RNAcentral gene coordinate entry. Differential count statistics corresponding to small RNA changes were derived from the exactTest function of the edgeR package for differential expression analysis^83,84^ over the entire RNAcentral annotation set. Downstream analyses and visualizations subset on canonical Pol III-transcribed genes and, specifically, the 350 genes confidently identified as Pol III-occupied in our system by ChIP-seq analysis.

### Correlation between Pol III subunit expression and gene chromatin features

[A] *Primary Immune cell populations*: Simultaneous ATAC-seq and gene expression data derived from primary immune progenitor and primary differentiated immune cell populations were obtained from GEO series GSE74246, GSE74912, GSE118165, and GSE118119^63,64^. Pol III subunit expression profiles were extracted from previously processed gene abundance analysis files (GSE74246_RNAseq_All_counts.txt and GSE118165_RNA_gene _abundance.txt). For the purpose of Pol III identity and gene accessibility correlations, the log2 ratio of POLR3G and POLR3GL abundance levels within individual samples was considered. Raw ATAC-seq data were downloaded and mapped to the GRCh38 genome using bowtie version 2.2.4 with settings “bowtie2 -t –sensitive -x”^82^. Read counts were extracted over a comprehensive RNAcentral database annotation of noncoding RNAs^54,55^. Gene annotation set and corresponding GRCh38 coordinates were downloaded from the contemporaneous RNAcentral database (release version13): ftp://ftp.ebi.ac.uk/pub/databases/RNAcentral/releases/13.0/genome_coordinates/bed/homo_sapie ns.GRCh38.bed.gz. Gene-specific raw ATAC-seq signal counts were extracted and include the flanking 150 bp for each RNAcentral gene coordinate entry. Gene accessibility signal counts were rank normalized across 248 ATAC-seq experiments, including unstimulated and stimulated immune cell experiments^64^. In total, 188 simultaneous ATAC-seq and RNA-seq profiles were matched by sample ID, and the Pearson correlation determined for the log2(POLR3G/POLR3GL) ratio and normalized gene accessibility profile in individual samples (Extended Data Fig. 3d). Analogous Pol III identity and gene chromatin feature correlation analysis was performed against 10 aggregate cell populations, including progenitor, myeloid, B cells (unstimulated), natural killer (NK) cells (unstimulated), CD4+ T cells (unstimulated), CD8+ T cells (unstimulated), and γδ T cells (unstimulated), pre-leukemic hematopoietic stem cells (pHSCs), leukemia stem cells (LSC), and leukemic blast cells (Blast) (Extended Data Fig. 3d).

[B] *The Cancer Genome Atlas* (TCGA): Pol III subunit expression levels were extracted across all TCGA disease cohorts from the Broad Institute TCGA Genome Data Analysis Center (GDAC) Firehose mRNASeq Level 3 RSEM gene normalized data files at https://gdac.broadinstitute.org. For analysis of Pol III subunit expression changes in individual disease cohorts, patient-matched normal tissue (11) and primary solid tumor (01) expression profiles were merged by participant ID. Disease cohorts with >= 5 patient-matched samples were considered (BLCA, n=19; BRCA, n = 115; CHOL, n=9; COADREAD, n=32; ESCA, n=11; KICH, n=25; LUAD, n=58; LUSC, n=51; PRAD, n=52; STAD, n=32; THCA, n=59; UCEC, n=10). Primary solid tumor ATAC-seq alignment bam files were retrieved from the Genomic Data Commons Data Portal (https://portal.gdc.cancer.gov). Read counts were extracted over a comprehensive RNAcentral database annotation of noncoding RNAs^54,55^. Gene annotation set and corresponding GRCh38 coordinates were downloaded from the contemporaneous RNAcentral database (release version13): ftp://ftp.ebi.ac.uk/pub/databases/RNAcentral/releases/13.0/genome_coordinates/bed/homo_sapiens.GRCh38.bed.gz. Gene-specific raw ATAC-seq signal counts were extracted and include the flanking 150 bp for each RNAcentral gene coordinate entry. Gene accessibility signal counts were rank normalized across 410 TCGA ATAC-seq experiments. In total, 388 ATAC-seq and RNA-seq profiles were matched by primary solid sample ID, and the Pearson correlation determined for the log2(POLR3G/POLR3GL) ratio and normalized gene accessibility profile in individual samples (Extended Data Fig. 4d).

### Multi-experiment overlapping gene enrichment analysis

Moving overlap analysis identifies the number of genes, when ranked by degree of gene loss (e.g. Pol III occupancy, small RNA abundance +/− PMA; ML-60218), that are shared between 2 or more independent experiments (e.g. dynamic Pol III occupancy v. dynamic small RNA abundance +/− PMA; PMA-sensitive v. ML-60218 sensitive RNA abundance, etc.) at a given cutoff level wherein the number of top genes (n) are considered. Maximum enrichment scores (Fig. 5p) represent the highest observed/expected ratio for a given gene based on representation within an overlapping gene set along the moving plot.

## DECLARATIONS

## Acknowledgements

We thank Dr. Joshua J. Gruber, Dr. E. Peter Geiduschek, Dr. Eric Phizicky, Dr. Howard Chang, Dr. Rebecca Harper, and members of the Kornberg and Snyder lab for insightful discussion and feedback. Illumina sequencing services were performed by the Genome Sequencing Service Center by Stanford Center for Genomics and Personalized Medicine Sequencing Center. The results in this study are in part based upon data generated by the TCGA Research Network: https://www.cancer.gov/tcga

## Availability of data and materials

The sequencing data from this study have been deposited through the NCBI Gene Expression Omnibus (GEO; http://www.ncbi.nlm.nih.gov/geo/) under GEO series number GSE163422 and series number GSE171884, and are publicly available.

## Ethics approval and consent to participate

Not applicable

## EXTENDED DATA FIGURES

**Extended Data Fig. 1.**
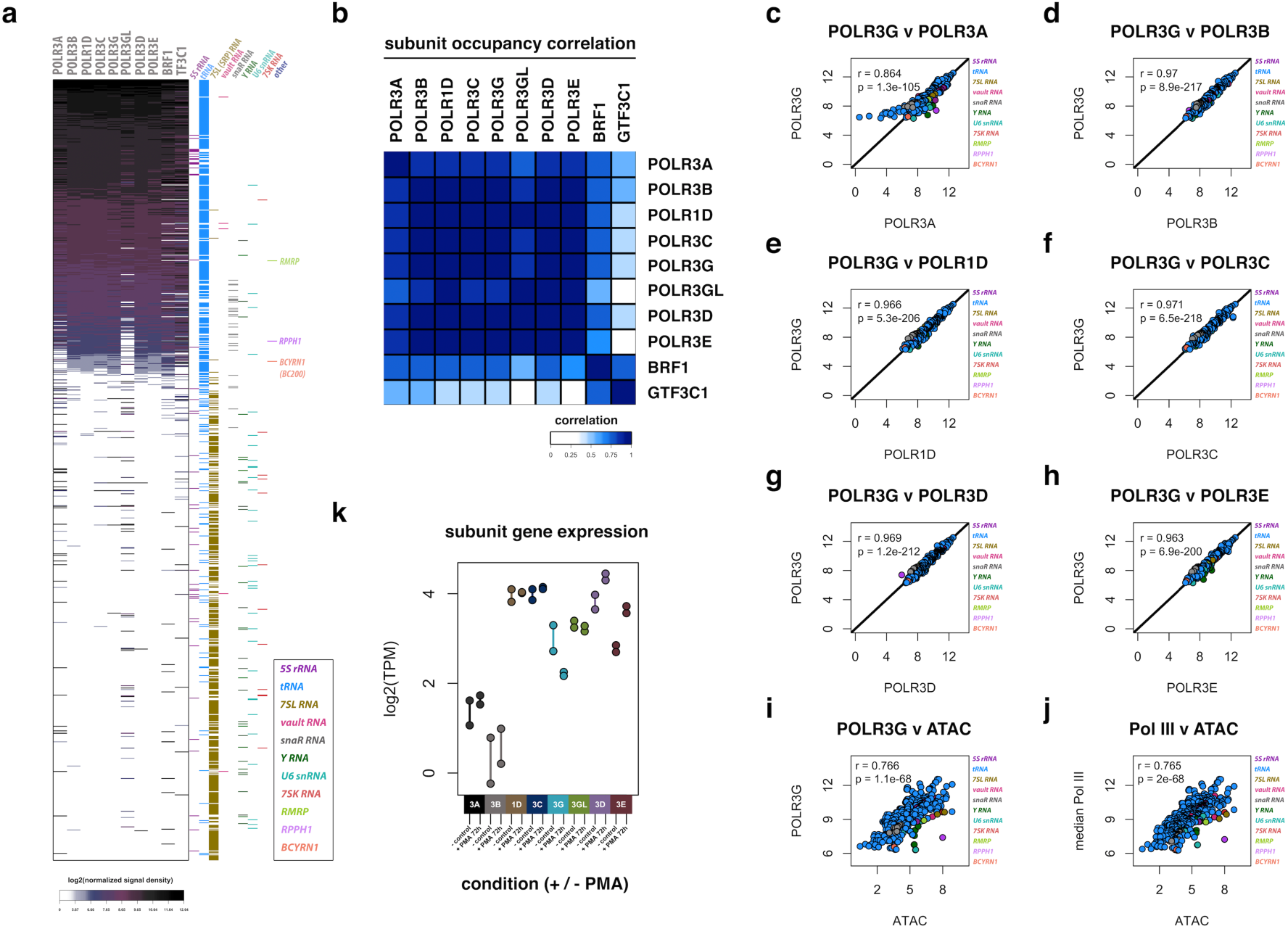
Pol III subunit expression, genomic occupancy, and chromatin accessibility. (**a**) Heatmap visualization of ChIP-seq signal density over individual Pol III-transcribed genes for Pol III subunits POLR3A, POLR3B, POLR1D, POLR3C, POLR3G, POLR3GL, POLR3D, and POLR3E, TFIIIB subunit BRF1, and TFIIIC subunit TF3C1. Heatmap is ordered by the median signal density across Pol III subunits over canonical Pol III-transcribed genes (n = top 1,000 canonical genes; RNAcentral annotation). Corresponding gene type indicated by right-flanking colorbar. (**b**) Heatmap visualization of Pearson correlation between chromatin IP experiments for individual Pol III subunits and BRF1, GTF3C1. (**c-h**) Correlation scatterplot visualization between Pol III subunit POLR3G and Pol III subunits POLR3A, POLR3B, POLR1D, POLR3C, POLR3D, and POLR3E, respectively. (**i**) Scatterplot visualization of correlation between Pol III subunit POLR3G signal enrichment and chromatin accessibility profile (ATAC-seq) at Pol III occupied genes in THP-1 monocytes. (**j**) Analogous correlation plot between chromatin accessibility profile (ATAC-seq) and median Pol III subunit signal enrichment. (**k**) Gene expression profile for mapped Pol III subunits in THP-1 cells (points represent individual biological replicates).

**Extended Data Fig. 2.**
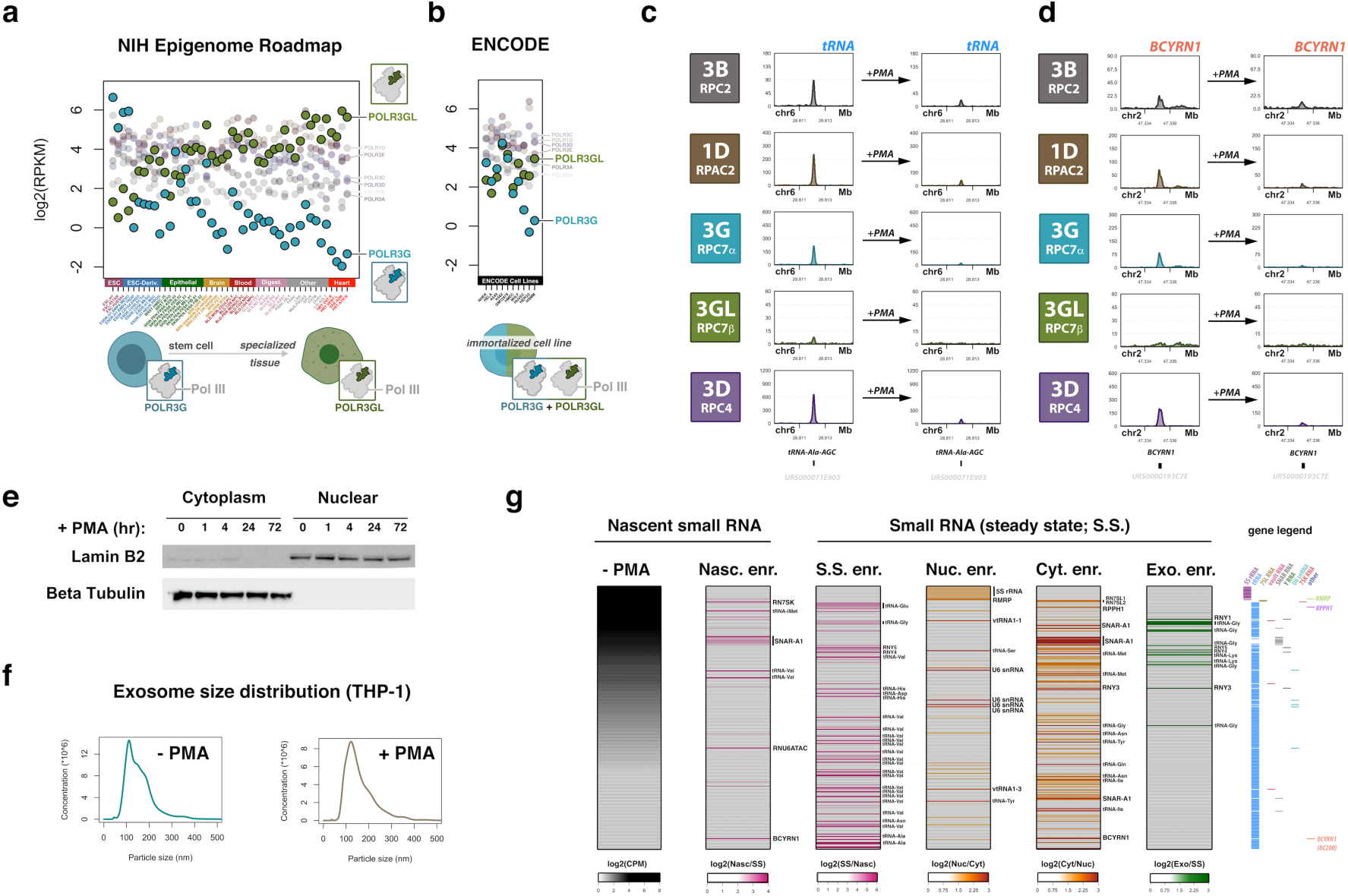
Dynamic Pol III subunit expression, occupancy, and compartmental small RNA abundance in THP-1. (**a**) Gene expression levels of mapped Pol III subunits across distinct cellular and tissue contexts with emphasis on POLR3G and POLR3GL expression in embryonic stem cells (ESCs), ESC-derived, epithelial, brain, blood, digestive, heart, and other contexts (NIH epigenome roadmap). (**b**) Gene expression levels of mapped Pol III subunits across immortalized ENCODE cell lines. (**c**) ChIP-seq profile for POLR3B, POLR1D, POLR3G, POLR3GL, and POLR3D are shown in THP-1 cells before (left) and after 72 h PMA treatment (right) over tRNA gene *Ala-AGC* (URS000071E903). (**d**) Analogous ChIP-seq signal profile over *BCYRN1* gene encoding BC200 RNA before and after 72 h PMA treatment in THP-1 cells. (**e**) Subcellular fractionation time course immunoblot for Lamin B2 (nuclear marker), and Beta Tubulin (cytoplasm marker) protein levels in THP-1 cells at 0, 1, 4, 24, and 72 h post PMA treatment. Immunoblot represents subcellular fractionation experiments corresponding to nuclear and cytoplasmic small RNA purification. (**f**) Purified THP-1 exosome nanoparticle size analysis. THP-1 monocyte and THP-1 macrophage exosomes were isolated by differential centrifugation and exosome particle size distributions were assessed by nanoparticle tracking analysis (NTA; NanoSight). Plots represent NTA of particle concentration (particle/mL) and size distribution of THP-1 monocyte exosomes (left) and THP-1 macrophage exosomes (right). (**g**) Heatmap visualization of small RNA profiles in THP-1 monocytes, including analyses of nascent, steady-state, nuclear, cytoplasmic, and exosomal small RNA enrichment. All heatmaps are ordered by the level of nascent RNA abundance on far left heatmap. Gene legend for all heatmaps on right – rows include all canonical Pol III-transcribed genes occupied by Pol III subunits in THP-1. From left-to-right, heatmaps visualize nascent RNA abundance (- PMA), nascent enrichment over steady-state abundance (Nasc. enr.), steady-state enrichment over nascent abundance (S.S. enr.), nuclear subcellular fraction enrichment over cytoplasmic fraction (Nuc. enr.), cytoplasmic subcellular fraction enrichment over nuclear fraction (Cyt. enr.), and exosomal purified small RNA enrichment over total cellular abundance (Exo. Enr.)

**Extended Data Fig. 3.**
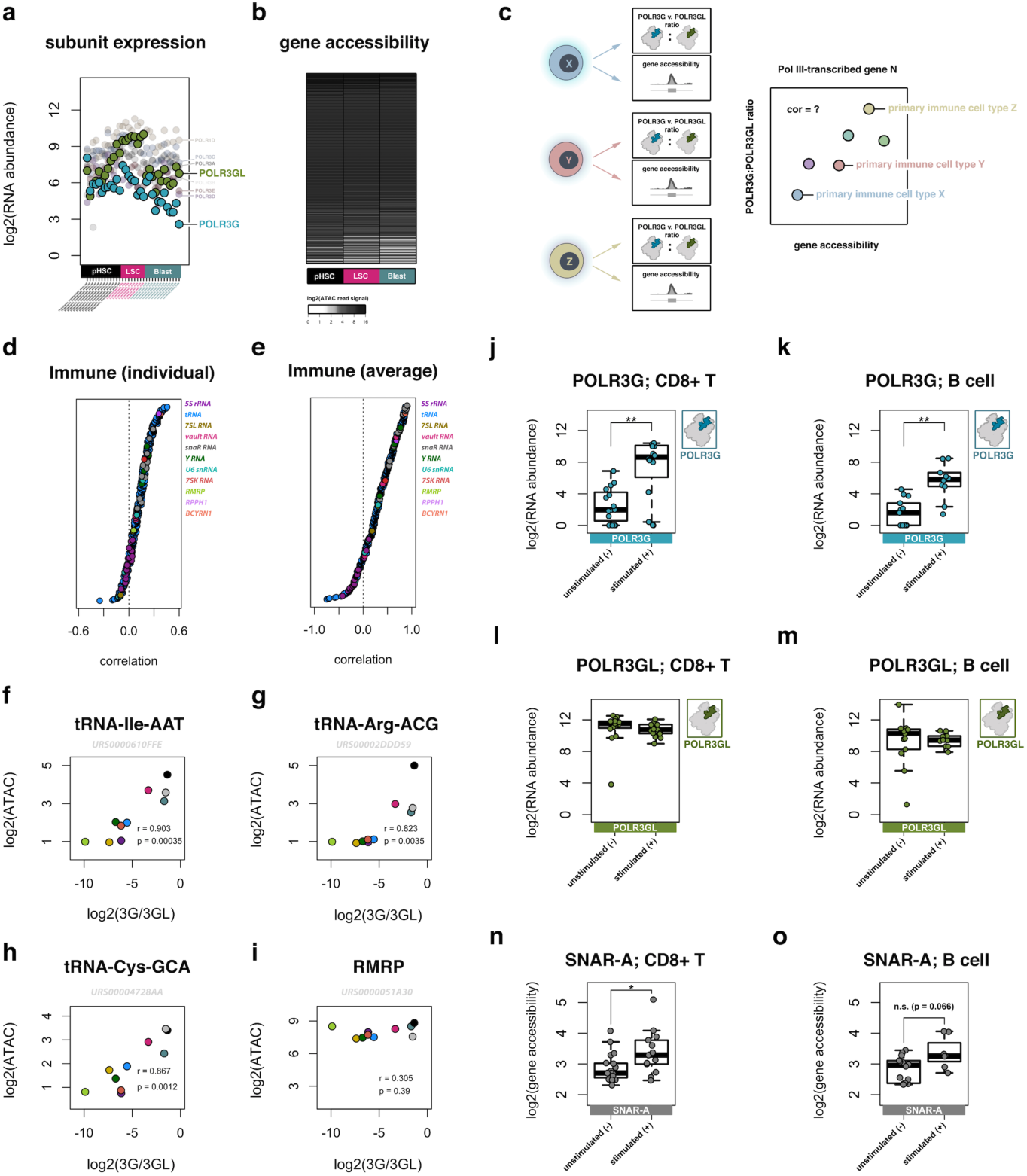
Integrated analysis of Pol III identity and transcriptome chromatin features during primary immune differentiation. (**a**) Gene expression levels of mapped Pol III subunits across pre-leukemic hematopoietic stem cells (pHSC), leukemia stem cells (LSC), and Leukemia blast cells (Blast) with emphasis on POLR3G and POLR3GL. (**b**) Gene accessibility measured by ATAC-seq read signal mapping to the repertoire of Pol III occupied genes corresponding to gene expression profiles and cell types in Extended Data Fig. 3a. (**c**) Illustration of approach for integrated analysis of Pol III identity and gene repertoire activity in primary immune cells. POLR3G and POLR3GL expression ratios and ATAC-seq accessibility signals are matched to individual samples and the relationship between these features are assessed by correlation analysis for each individual Pol III-transcribed gene across all profiles samples. (**d**) Resulting correlation scores for Pol III-transcribed genes across all individual samples. (**e**) Resulting correlation scores for Pol III-transcribed genes across aggregate cell-type populations. (**f-i**) Example profiles of individual gene accessibility correlation scores with POLR3G:POLR3GL expression ratios in primary immune cells, including (**f**) *tRNA-Ile-AAT* (URS0000610FFE) (**g**) *tRNA-Arg-ACG* (URS00002DDD59) (**h**) *tRNA-Cys-GCA* (URS00004728AA) and (**i**) *RMRP*. (**j-k**) POLR3G expression in primary CD8+ T cells and primary B cells before and after stimulation (anti-CD3/CD28 and IL-4 respectively) (**l-m**) POLR3GL expression in primary CD8+ T cells and primary B cells before and after stimulation (**i**) *SNAR-A* gene accessibility profiles as measured by ATAC-seq in primary CD8+ T cells and primary B cells before and after stimulation.

**Extended Data Fig. 4.**
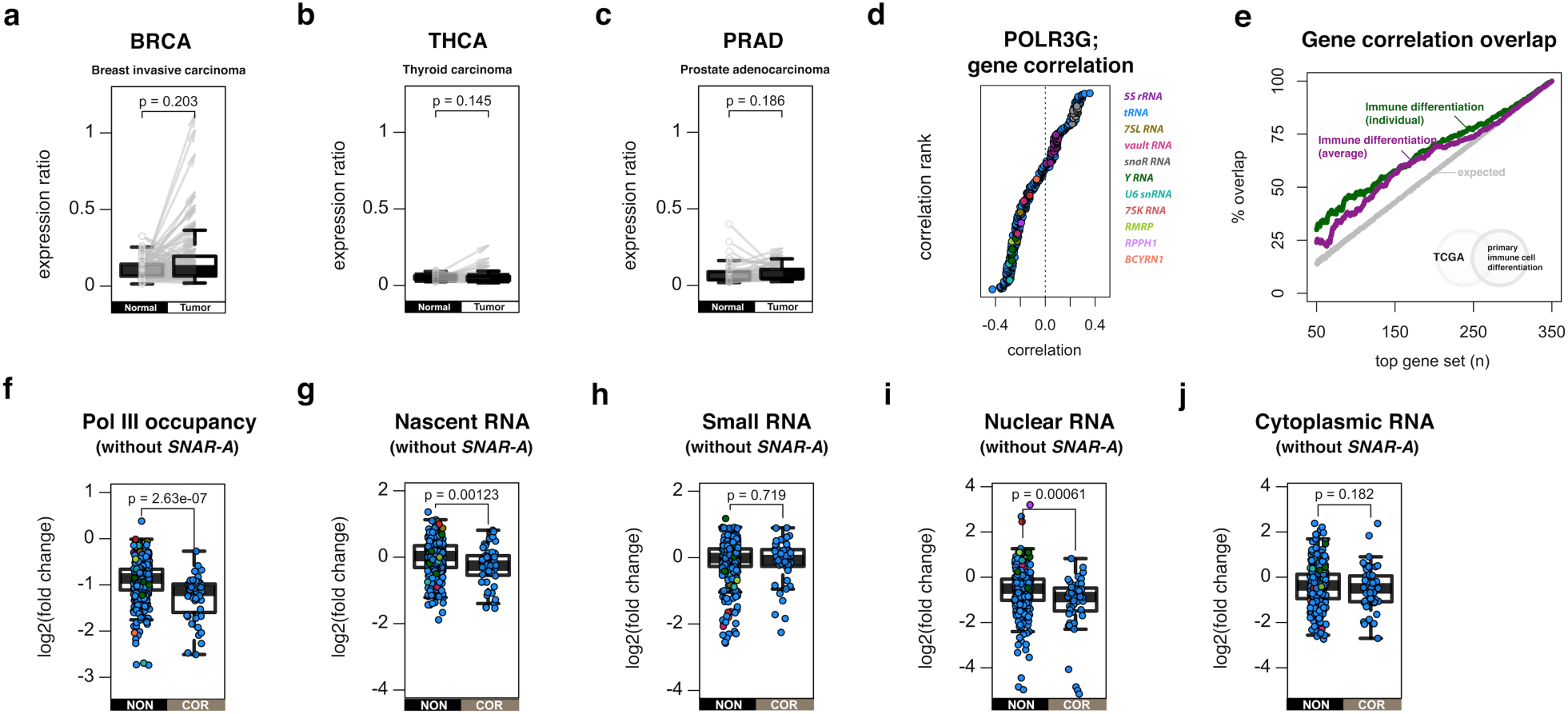
POLR3G expression and correlative chromatin signatures at POLR3G-sensitive gene repertoires in human primary solid tumors. (**a-c**) Gene expression ratios of POLR3G/POLR3GL in matched normal tissue and primary solid tumors in breast invasive carcinoma (**a**, n=115), thyroid carcinoma (**b**, n=59), and prostate adenocarcinoma (**c**, n=52) primary solid tumors. (**d**) Ranked correlation scores for Pol III-transcribed gene accessibility and POLR3G/POLR3GL ratios across individual primary solid tumors. Points represent individual Pol III-transcribed genes; gene type indicated by color legend. (**e**) Moving Venn-diagram overlap analysis of the strongest positively correlating genes in primary immune cells (individual sample analysis = green; aggregate cell type analysis = purple) with gene correlation scores in TCGA primary solid tumors. (**f-j**) Comparison analysis of changes in correlative (COR) and noncorrelative (NON) subgroups (with pre-removal of *SNAR-A* genes) related to Pol III occupancy (**f**), nascent RNA (**g**), steady-state RNA (**h**), nuclear (**i**), and cytoplasmic RNA (**j**) small RNA in THP-1 cells +/− 72h PMA-induced differentiation and entry to a quiescent cellular state.

**Extended Data Fig. 5.**
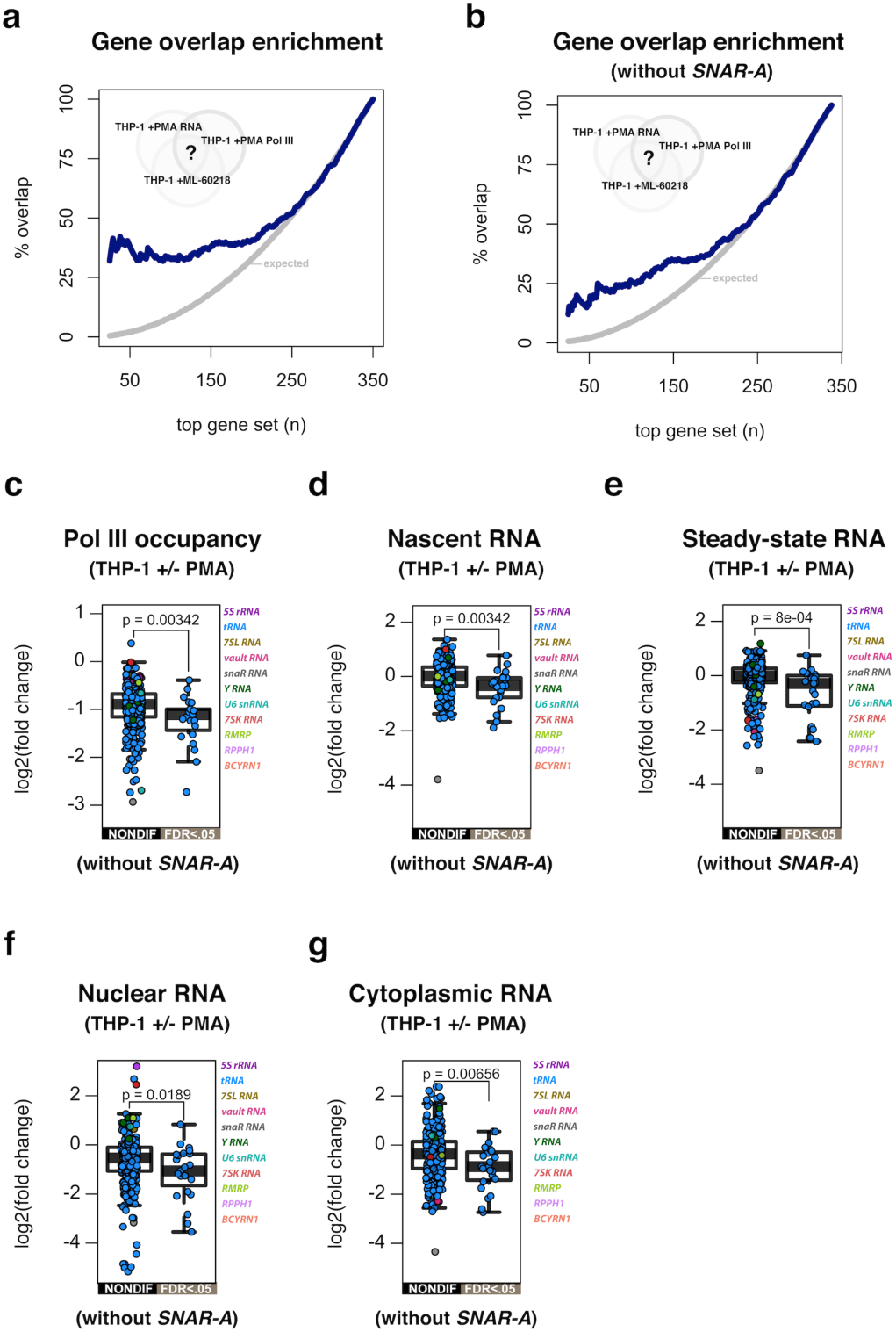
POLR3G disruption mirrors dynamic RNA signatures observed during cell differentiation. (**a**) Moving Venn-diagram overlap analysis of the dynamic Pol III occupancy and small RNA response observed during THP-1 differentiation (72 h PMA exposure) with POLR3G disruption effects on THP-1 small RNA levels (4 h ML-60218 exposure). (**b**) Analogous overlap enrichment analysis of the dynamic Pol III occupancy and small RNA response observed during THP-1 differentiation (72 h PMA exposure) with POLR3G disruption effects on THP-1 small RNA levels (4 h ML-60218 exposure) excluding *SNAR-A* gene representation. (**c-g**) Comparison of the log2(fold change) in Pol III occupancy (**c**), nascent RNA levels (**d**), total steady-state small RNA levels (**e**), nuclear RNA levels (**f**), and cytoplasmic RNA levels (**g**) for nondifferential (NONDIF) and significantly differential (FDR<.05) genes that are sensitive to 4-hour ML-60218 exposure in THP-1 cells, before and after PMA-induced differentiation, excluding *SNAR-A* genes (related to Fig. 5j-n).

**Extended Data Fig. 6.**
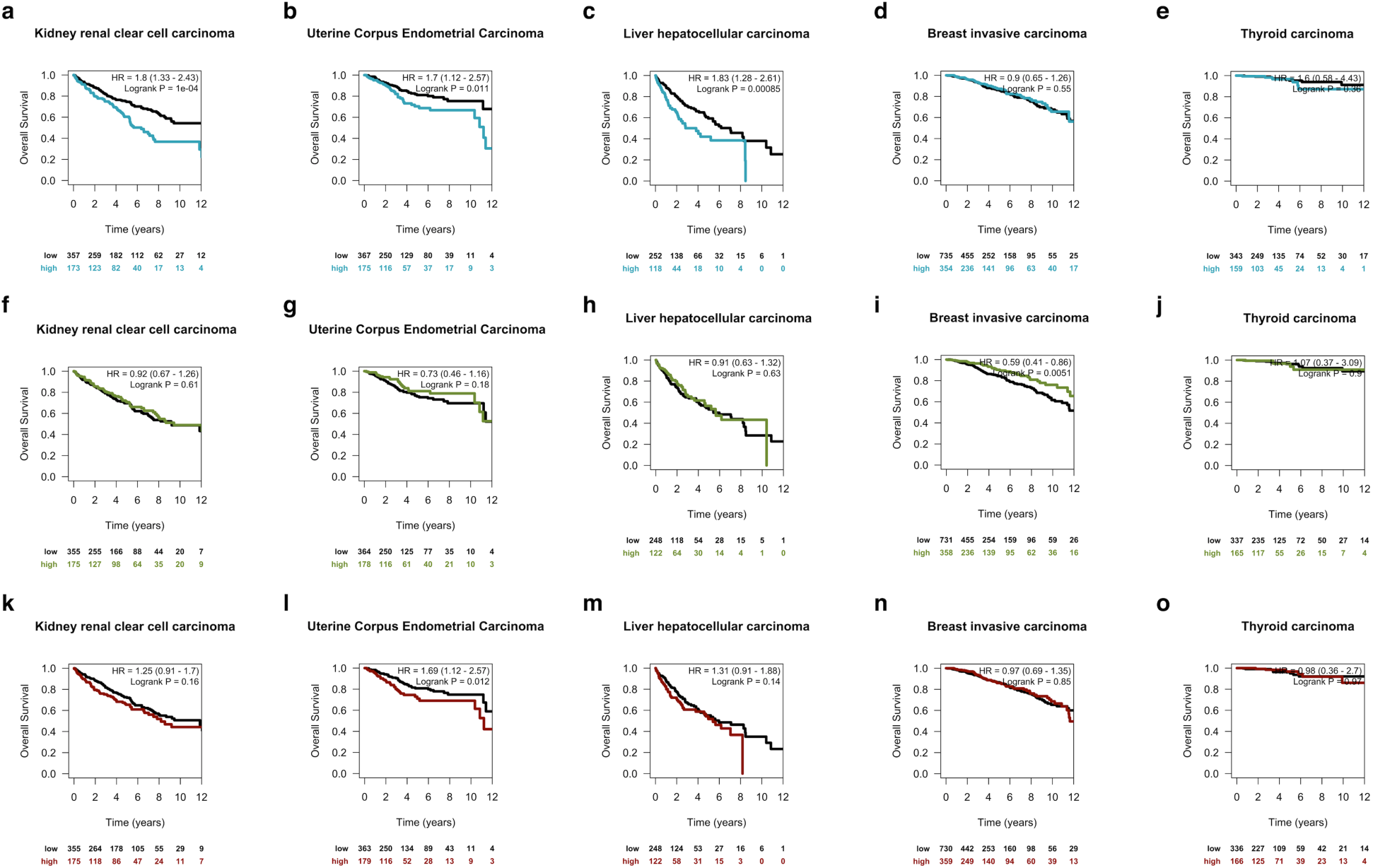
POLR3G levels are associated with poor survival outcomes in specific cancers. (**a-e**) Kaplan-Meier analysis of overall survival of TCGA donors stratified by high POLR3G expression (top tertile) and normal POLR3G expression in kidney renal clear cell carcinoma (**a**), uterine corpus endometrial carcinoma (**b**), liver hepatocellular carcinoma (**c**), breast invasive carcinoma (**d**), and thyroid carcinoma (**e**). (**f-j**) Kaplan-Meier analysis of overall survival of TCGA donors stratified by high POLR3GL expression (top tertile) and normal POLR3GL expression in kidney renal clear cell carcinoma (**f**), uterine corpus endometrial carcinoma (**g**), liver hepatocellular carcinoma (**h**), breast invasive carcinoma (**i**), and thyroid carcinoma (**j**). (**k-o**) Kaplan-Meier analysis of overall survival of TCGA donors stratified by high MYC expression (top tertile) and normal MYC expression in kidney renal clear cell carcinoma (**k**), uterine corpus endometrial carcinoma (**l**), liver hepatocellular carcinoma (**m**), breast invasive carcinoma (**n**), and thyroid carcinoma (**o**). HR = hazard ratio risk of dying.

